# Systemic delivery of drug-free polymeric nanoparticles reprograms innate immunity in a sex-dependent manner after spinal cord injury

**DOI:** 10.64898/2026.03.06.709912

**Authors:** Jaechang Kim, Irina Kalashnikova, Ruby Maharjan, Fernanda Stapenhorst França, Daniel Kolpek, James Ogidi, John C. Gensel, Jonghyuck Park

## Abstract

Sex differences influence distinct inflammatory responses after spinal cord injury (SCI), yet their impact on immune-modulating nanotherapeutics remains unclear. Here, we investigated the sex-dependent effects of drug-free poly(lactic-co-glycolic acid) (PLGA)-based nanoparticles (NPs) following SCI. Systemic NP administration enhanced locomotor recovery in both sexes and eliminated the functional gap observed in controls. Mechanistically, NPs engaged distinct immune pathways between sexes. Females accumulated more NPs in the spleen, leading to reduced monocyte-derived macrophage infiltration, whereas males showed greater NP accumulation at the lesion and attenuated microglial activation. Transcriptomic analysis showed preferential modulation of eicosanoid-related pathways in females and NF-κB-linked signaling in males. These sex-specific, yet convergent NPs-induced immunomodulatory effects reduced fibrotic scarring and enhanced remyelination, with females showing greater Schwann cell-mediated repair and males exhibiting marked suppression of microglial activation. Collectively, these findings demonstrate that NPs promote comparable functional recovery in both sexes through distinct, sex-influenced immune mechanisms and establish a translational framework for sex-informed immune targeting and nanotherapeutic design in SCI and other inflammation-mediated diseases.

**Graphic Abstract:** 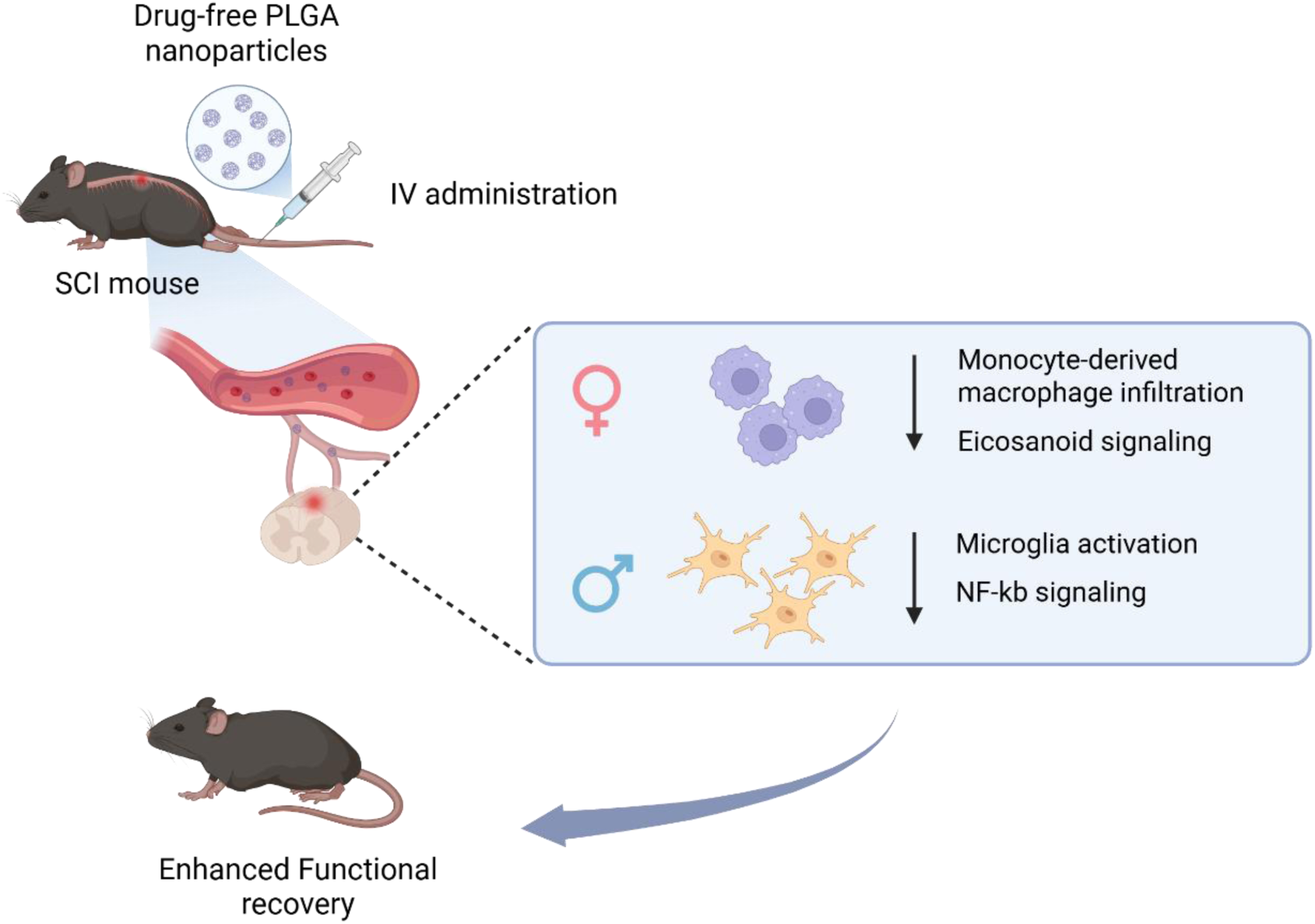

## Introduction

Spinal cord injury (SCI) is a devastating condition that results from traumatic events such as falls or motor vehicle accidents and often leads to permanent motor, sensory, and autonomic dysfunction [1, 2]. Beyond the traumatic primary injury, a prolonged secondary injury driven largely by inflammation promotes neuronal cell death, demyelination, and functional deficits [3, 4]. Within hours to days after injury, circulating innate immune cells such as neutrophils, inflammatory monocytes, and macrophages infiltrate the lesion site through the compromised blood–spinal cord barrier, initiating a robust inflammatory response [4, 5]. Although these cells aid in debris clearance and secrete growth-promoting factors, they also release proteases, reactive oxygen species (ROS), and multiple pro-inflammatory cytokines that exacerbate tissue damage [6, 7].

Because inflammation plays a central role in secondary events, biological variables that modulate immune responses can critically influence therapeutic outcomes. In particular, sex has emerged as a key determinant of neuroinflammation after SCI [8, 9]. Males and females display distinct immune cell dynamics and transcriptional profiles following SCI, with males exhibiting stronger microglial activation and cytokine-related pathways, whereas females showing a higher proportion of monocyte-derived macrophages (MDMs) and upregulation of genes associated with DNA damage repair within the injured spinal cord [10, 11]. These sex-dependent inflammatory responses suggest that immunotherapies may act through fundamentally different cellular and molecular pathways in males and females.

Nanoparticles (NPs) have emerged as a promising strategy to modulate post-SCI inflammation [12, 13]. Polymeric NPs, including those made from poly(lactic-co-glycolic acid) (PLGA), can deliver therapeutic molecules and also exert intrinsic immunomodulatory effects without additional cargos. Notably, cargo-free PLGA-based NPs are readily taken up by circulating innate immune cells and modulate inflammatory responses through receptor-mediated interactions [14]. In particular, their biodegradation products, glycolic and lactic acid, can further regulate inflammatory signaling by limiting nuclear factor-kappa B (NF-κB) activation and influencing immune polarization [15, 16]. Our previous studies demonstrated that PLGA-based NPs interact with scavenger receptors particularly macrophage receptor with collagenous structure (MARCO) on circulating myeloid cells, influencing NPs uptake, biodistribution, and downstream immune responses after SCI [17]. MARCO-mediated internalization can redirect myeloid cell trafficking to the spleen and alter inflammatory gene expression, contributing to the therapeutic outcomes of drug-free PLGA NPs. We further demonstrated that systemic administration of these NPs reduces inflammatory cells accumulation and promotes pro-regenerative immune phenotype, improving both functional and anatomical recovery after SCI [17, 18].

Despite this therapeutic potential, sex differences in NP responsiveness remain largely unexplored. Most preclinical SCI studies using NPs have been conducted in females [9], yet sex-specific inflammatory responses may alter NP biodistribution, cellular uptake, and therapeutic impact. Evidence from traumatic brain injury (TBI) models support this concern, demonstrating sex-dependent variation in NPs accumulation and antioxidant NPs efficacy [19, 20]. These observations highlight a critical knowledge gap regarding how males and females may differentially respond to NP-based immunomodulation.

To address this gap, we investigated how male and female respond to systemic administration of drug-free PLGA NPs following SCI. We assessed NP biodistribution, innate immune cell recruitment, transcriptomic changes, tissue repair, and functional recovery to determine whether sex affects NP-mediated immunomodulation. By directly comparing responses across sexes, we aim to elucidate sex-associated determinants of NPs efficacy and to define whether PLGA NPs modulate the post-injury environment through distinct mechanisms in each sex. Understanding these interactions will be essential for the rational design of nanotherapeutic strategies that incorporate sex as a critical biological variable in future immune-modulating treatment, with relevance to a broad range of inflammation-mediated diseases.

## Materials and Methods

### Nanoparticle fabrication and characterization

Carboxylated PLGA (50:50) purchased from Nanosoft Polymers (Winston-Salem, NC) was dissolved in dichloromethane and emulsified in 2% poly(ethylene-alt-maleic anhydride) (PEMA) solution using a Qsonica Q125 Sonicator Ultrasonic Homogenizer W/Probe 125 W (ColeParmer; Vernon Hills, IL). The emulsion was poured into 0.5% PEMA solution and stirred at 400 rpm for the evaporation of the organic solvent. The suspension was washed three times and lyophilized with sucrose and D-mannitol (Sigma-Aldrich; St. Louis, MO). For the injections, NPs were resuspended in PBS at a concentration of 5mg/ml and filtered using a 0.45 µm mesh filter. The size, zeta potential, and polydispersity index were determined by using a Zetasizer Nano ZS (Malvern Panalytical; Malvern, UK) and the morphology was examined using FEI Quanta 250 field-emission scanning electron microscopy (FE-SEM).

### Spinal cord injury model and NPs administration

All procedures are approved by the University of Kentucky’s Institutional Animal Care and Use Committee. Female and male C57BL/6 mice (6-10 weeks old; Jackson Laboratories) were anesthetized using ketamine and xylazine, then received a laminectomy only as a sham injury, or laminectomy with 50- or 60-kDyn SCI at the T10 level using the Infinite Horizons Impactor (Precision Systems Instrumentation, LLC; Fairfax Station, VA). Mice analyzed for the chronic phase were subjected to a moderate SCI (60-kDyne) to induce sufficient deficits for distinguishing between control and treatment groups. The muscles were sutured together, then the skin was stapled. The animals were placed on a heating pad for recovery. All mice received buprenorphine (Buprenorphine SR, 1.0mg/kg day of surgery), as well as an antibiotic (Enrofloxacin, 8.5 mg/kg) and lactate ringer solution for up to 5 days post-SCI. Manual bladder expressions were performed 2x/day until bladder reflexive function was observed. Each animal received 50mg/kg of NPs (5mg/ml of suspension) via tail veil injection within two hours after SCI per day for 7 consecutive days. The same volume of PBS was injected as a control. Mice were excluded from the analysis if they experienced an improper injury or exhibited a BMS score greater than 3 at 1-day post injury (DPI).

### Basso mouse scale (BMS) for locomotion

The hindlimb movement of mice was evaluated using the BMS scoring system [21]. The scores of the left and right limbs were averaged to generate a single score for each mouse [22]. The BMS score ranges from 0 (complete paralysis) to 9 (normal movement) points, and the sub-score ranges from 0 to 11 points. Mice were observed in an open field for 4 minutes after they had gently adapted to the field. The scores are assessed on day 1 and 3 and then weekly after SCI for 8 weeks. The assessment was performed by researchers blinded to the group.

### Open-Field Spontaneous Activity Test

The open-field spontaneous activity test was used to measure locomotor activity at 56-DPI. Before testing, mice were acclimatized in the experimental room for 30 min [23]. Each mouse was placed individually in an open-field box (40 cm × 40 cm) for evaluation. The test parameters, including total distance traveled, average speed, and total time mobile, were recorded automatically for 5 min using an ANY-MAZE video tracking system (Stoelting Co.; Wood Dale, IL).

### Analysis of in vivo biodistribution

For the in vivo biodistribution, mice were administered NPs-Cy5.5 via the tail vein within two hours after SCI per day for 7 days. At 7-DPI, mice were euthanized, and the spinal cord and spleen were collected. Tissues were imaged using an Xenogen IVIS-200 imaging system (PerkinElmer; Waltham, MA) and analyzed by the Living Image Analysis Software.

### Flow cytometry analysis

Mice were euthanized, and a 4 mm section of the spinal cord with the center of the injury lesion was dissected and placed in a digestion buffer containing a 1:1 solution of Dulbecco’s Modified Eagle Medium (DMEM) and Accumax (Thermo Fisher Scientific; Waltham, MA). The cords were mashed through a 70 µm mesh filter into a 30% Percoll solution diluted in DMEM. Cells were centrifuged at 1500 g for 15 min at 10 °C to pellet cells, and the floating myelin layer was aspirated. Cells were then suspended in ACK lysing buffer (Quality Biological; Gaithersburg, MD) and pelleted. Cells were washed and suspended in flow cytometry staining buffer (Invitrogen; Carlsbad, CA) for antibody labeling. Endogenous Fc receptors were blocked with anti-CD16/32 (1:100; Cat #: 156604; BioLegend) antibody for 15 min. Cells were then incubated for 1 hour on ice with antibodies; Anti-CD45-PE/Cy7 (Cat #: 157206, BioLegend), Anti-CD11b-BV421 (Cat #: 101251, BioLegend), Anti-Ly6C-FITC (Cat #: 128006, BioLegend), Anti-Ly6G-PE (Cat #: 127608, BioLegend), Anti- Anti-CD206-APC (Cat #: 141708, BioLegend), Anti-CD163-PE (Cat #: 155307, BioLegend). The cells were analyzed using a flow cytometer (BD FACSymphony), and it was performed by the University of Kentucky’s Flow Cytometry and Immune Monitoring Facility.

### Tissue processing and Immunofluorescence

Mice were anesthetized using ketamine and xylazine and were perfused using cardiac puncture with PBS, followed by 4% paraformaldehyde in PBS. The spinal cords were extracted, then post-fixed for 2 hours at room temperature and transferred to a 30% sucrose solution at 4 °C before blocking in Tissue-Tek OCT compound (Sakura Finetek; Torrance, CA). Samples were sectioned in 18 µm transversely on an Epredia HM525 NX Cryostat.

Samples were washed with tris(hydroxymethyl)aminomethane (Tris) buffer solution, followed by permeabilization with Triton-X and blocking with 10% normal donkey serum solution at room temperature. Then they were stained with primary antibodies at 4°C and subsequently stained with secondary antibodies at room temperature. The detailed information about primary and secondary antibodies was provided in Table S1. Samples were washed with Tris buffer solution and incubated with Hoechst 33342 to stain nuclei. Images were acquired with a Nikon AXR inverted confocal microscope and analyzed using ImageJ.

### Gene expression using NanoString and RT-qPCR

Mice were euthanized, and the spinal cord of 4mm length centered on the injury site was homogenized using 1 ml of Trizol reagent (Invitrogen; Carlsbad, CA) with a glass tissue grinder. RNA was extracted according to the manufacturer’s instructions. For NanoString analysis, the concentration and quality of RNA were measured using the Epoch plate reader (BioTek; Winooski, VT) and BioAnalyzer (Agilent Technologies; Santa Clara, CA), respectively. RNA analysis was performed by NanoString nCounter Mouse Inflammation v2 Panel following the manufacturer’s instructions (NanoString Technologies; Seattle, WA) at the University of Kentucky’s Genomics Core Laboratory. Data was analyzed using nSolver software and visualized using R and GraphPad Prism.

For RT-qPCR, the concentration of RNA was measured as previously mentioned, and cDNA was synthesized using the iScript™ cDNA Synthesis kit (Bio-Rad; Hercules, CA). The RT-qPCR products were assessed using the accumulation level of iQ™ SYBR Green Supermix (Bio-Rad) fluorescence following a manufacturer’s protocol on CFX Connect™ Real-Time PCR Detection System (Bio-Rad). 18s-rRNA served as an internal reference gene for data normalization. The relative expression levels of the target genes were calculated using the ΔΔCt method. Detailed sequences of the primers used are listed in Table S2.

### Statistical analysis

Data are presented as means ± SEM and analyzed using GraphPad Prism software. Statistical differences between two groups were analyzed using an unpaired t-test, while differences among more than two groups were assessed using two-way analysis of variance (ANOVA) with Tukey’s multiple post hoc comparisons. Statistical significance was considered at p <0.05 (*), p <0.01 (**), p<0.001 (***), p<0.0001 (****).

## Results

### PLGA NPs promote locomotor functional recovery and reduce sex-based differences after SCI

We previously demonstrated that PLGA NPs enhance functional recovery in females after SCI [17, 18]. In this study, we compared the functional recovery between sexes following NP treatment. The NPs possessed a negative surface charge and an average diameter of approximately 500 nm (Fig. 1A). NPs administered intravenously at 50 mg/kg once daily for 7 days, beginning within 2 hours after T10 contusion SCI (Fig 1B), aligning with the acute phase of innate immune cells infiltration that begins shortly after injury [17].

**Figure 1.**
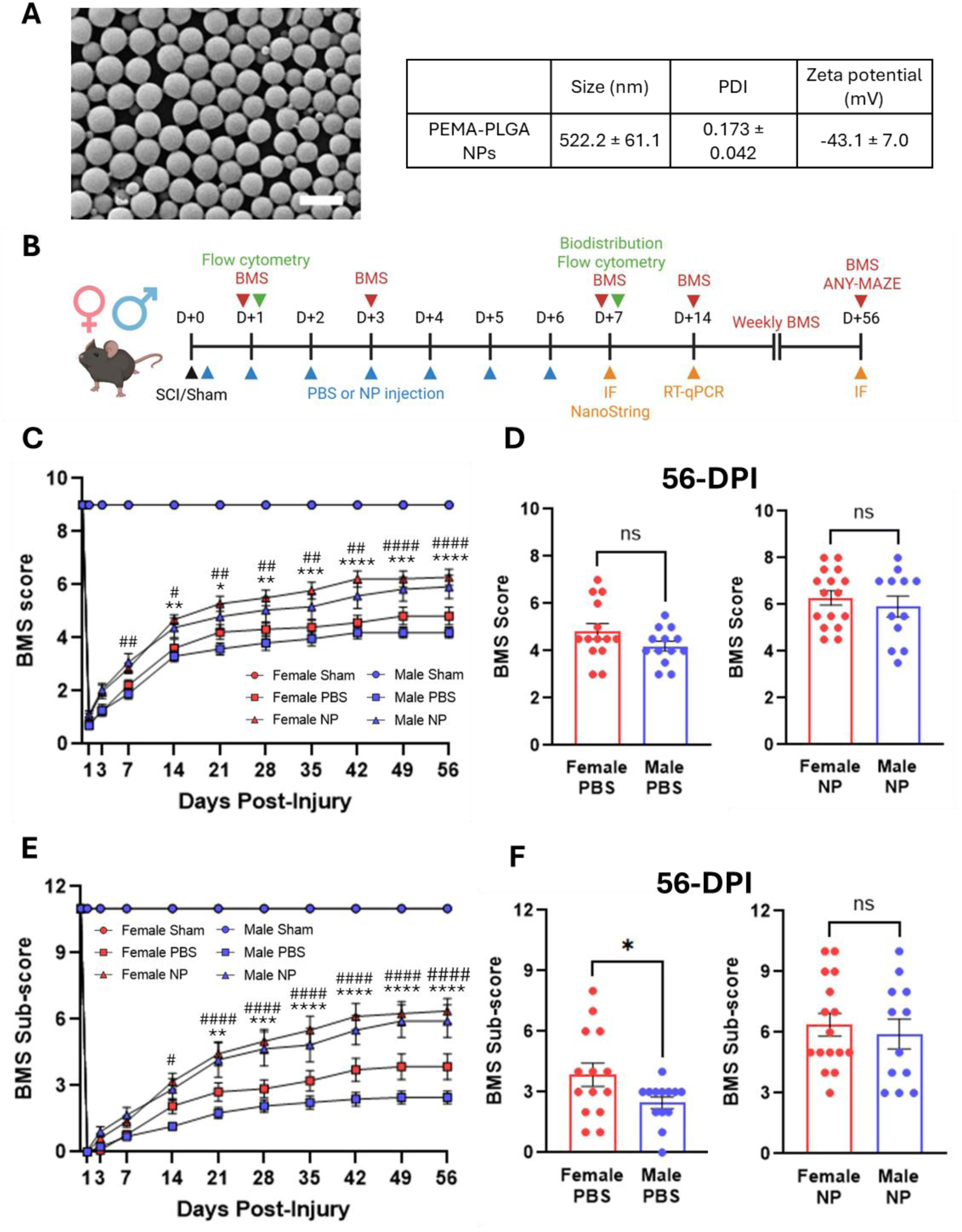
PLGA NPs promoted locomotor function recovery and reduced the sex-dependent differences. (A) Morphology and physicochemical characterization of PLGA NPs. Scale bar = 1 µm. (B) Schematics of the experimental workflow. Created in BioRender. Nano, B. (2026) https://BioRender.com/58k2u8k. (C) The hindlimb locomotor recovery was assessed using the BMS weekly for 56-DPI. The locomotor function improved after NP treatment in both sexes. (D) Individual BMS scores of PBS and NP-treated groups at 56-DPI showed no significant difference between males and females. (E) The BMS sub-score was used to observe subtle locomotor recovery. The sub-score increased after NP treatment in both sexes. (F) Individual BMS sub-score of PBS and NP-treated groups in both sexes at 56-DPI. In the PBS groups, females had significantly higher sub-scores compared to males, but there was no difference in the NP-treated groups. For all test, male sham: n = 12; female sham: n = 10; male PBS: n = 13; female PBS: n = 14; male NP: n = 12; female NP: n = 16. For weekly BMS score and sub-score data, A two-way ANOVA with Tukey’s post hoc test for multiple comparisons. * p<0.05, ** p<0.01, *** p<0.001, and **** p<0.0001 compared with the PBS group in female, # p<0.05, ## p<0.01, and #### p<0.0001 compared with the PBS group in male. For BMS and sub-score data at 56-DPI, an unpaired t-test for the comparison. ns: no significant; * p < 0.05. Data are presented as mean ± SEM.

Hindlimb locomotor recovery was assessed using the BMS [21] for 56-DPI. Injury severity and body weight changes were comparable across groups (Fig. S1). All animals exhibited normal open field locomotion before SCI (BMS = 9). NP-treated mice exhibited markedly improved BMS scores relative to PBS controls in both sexes (Fig. 1C). In females, improvement emerged by 14-DPI and persisted through 56-DPI, whereas males showed significant benefit as early as 7-DPI. Notably, BMS scores did not differ between sexes within either treatment group (Fig. 1D).

Because specific components of locomotor recovery are not fully captured by the overall BMS score [21, 24], we also performed BMS sub-scoring in addition to the BMS to assess specific motor improvements such as stepping frequency, paw position, and trunk stability [25, 26] (Fig. 1E). PBS-treated females showed significantly higher sub-scores than PBS-treated males at 56-DPI (p<0.05; Fig. 1F). Notably, NP-treated males and females reached comparable sub-score, indicating that NP treatment mitigates the sex-related differences in locomotor recovery. Detailed behavioral data showing BMS score and sub-score distributions are shown in Supplementary Fig. 2.

To further evaluate spontaneous motor activity, mice were analyzed in an open-field chamber at 56-DPI using the ANY-MAZE tracking system [23], and it supported consistently with improved function after NP treatment. Although there was no significant difference between the PBS and NP-treated groups, the total distance traveled, average speed, and total time mobile tended to increase in the NP-treated groups compared to the PBS groups in both sexes (Fig. S3A). The images displaying locomotor traces of mice also demonstrated improved locomotor activity in NP-treated groups compared to PBS controls (Fig. S3B and S3C).

### Sex-Dependent Biodistribution of PLGA NPs in the spinal cord and spleen

To investigate the basis of the sex-dependent functional effects of NPs, we next assessed NPs’ in vivo biodistribution in male and female SCI mice. Cyanine 5.5 conjugated NPs (NPs-Cy5.5) were administered intravenously for 7 days, and the spinal cord and spleen were collected one day after the last administration. The fluorescence of NPs was readily detected at the contusion lesion in injured spinal cords but was absent in the sham groups (Fig. 2A). This indicates that acute disruption of the blood-spinal cord barrier after contusion allows NPs entry into the lesion site [27]. Radian efficiency levels associated with NPs-Cy5.5 in the spinal cord were significantly greater in the SCI groups compared to the sham groups in both sexes. However, in the spleen, no differences were observed between the sham and SCI groups (Fig. 2B).

**Figure 2.**
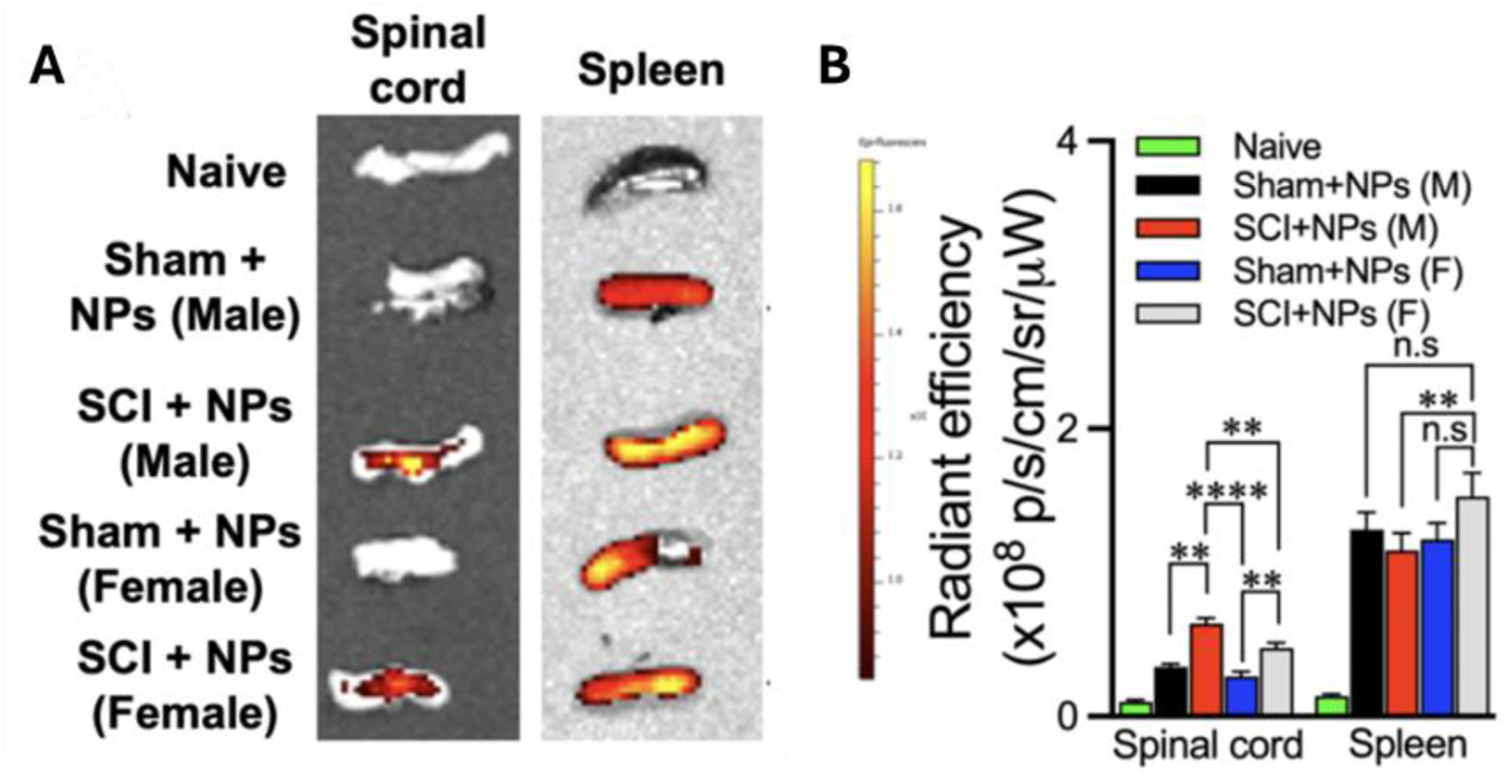
Intravenous administration of PLGA NPs has sex-dependent biodistribution. (A) Representative images of the spinal cord and spleen. NP accumulated in the spleen in all NP-treated groups, but in the spinal cord only in the SCI groups. (B) Quantification of fluorescence intensity. NP accumulation after SCI was higher in the spinal cord of males than females, whereas it was greater in the spleen of females than males. A two-way ANOVA with Tukey’s post hoc test for multiple comparisons. n = 5/group. n.s: no significant; ** p<0.01 and **** p<0.0001. Data are presented as mean ± SEM.

Sex-dependent differences in NP biodistribution were shown in comparison between the NP-treated males and females. Less accumulated NPs in the spinal cord were observed in the female group compared to the male group after SCI, which coincides with a substantial accumulation of NPs in the spleen in females (Fig. 2B; p<0.01 for each comparison). This sex-dependent pattern suggests enhanced sequestration of NP-associated circulating innate immune cells in females, leading to reduced NP accumulation within the injured spinal cord. These results indicate that systemic NP biodistribution after SCI is affected by sex, potentially associated with differences in innate immune cell uptake and trafficking between males and females.

### PLGA NPs exert sex-specific modulation of immune cell infiltration after SCI

To determine whether these biodistribution differences translate into distinct immune responses, we next analyzed immune cell populations in the injured spinal cord and spleen by flow cytometry at 1- and 7-DPI. The gating strategy is shown in Supplementary Fig. 4. These timepoints were chosen because the maximal infiltration day of neutrophils occurs at 1-DPI, and macrophages and microglia reach their peak at 7-DPI [27]. Flow cytometry analysis revealed significant sex differences in immune cell infiltration and response to NP treatment at 7-DPI. In the PBS groups, the proportion of infiltrated MDMs and microglia (CD45^+^/CD11b^+^/Ly6G^-^) was higher in females than in males (Fig. 3A), consistent with previous observations of increased MDM recruitment in females after SCI [11]. Remarkably, systemic NP administration substantially altered these patterns. In females, NPs significantly reduced the proportion of infiltrated MDMs and microglia compared to PBS (p<0.05), whereas males showed minimal change (Fig. 3A). Concomitantly, NP-treated females showed a significant increase in the splenic CD11b^+^ myeloid population (Fig. 3B), indicating enhanced sequestration of NP-associated myeloid cells in the spleen. In contrast, NPs did not significantly affect splenic myeloid accumulation in males.

**Figure 3.**
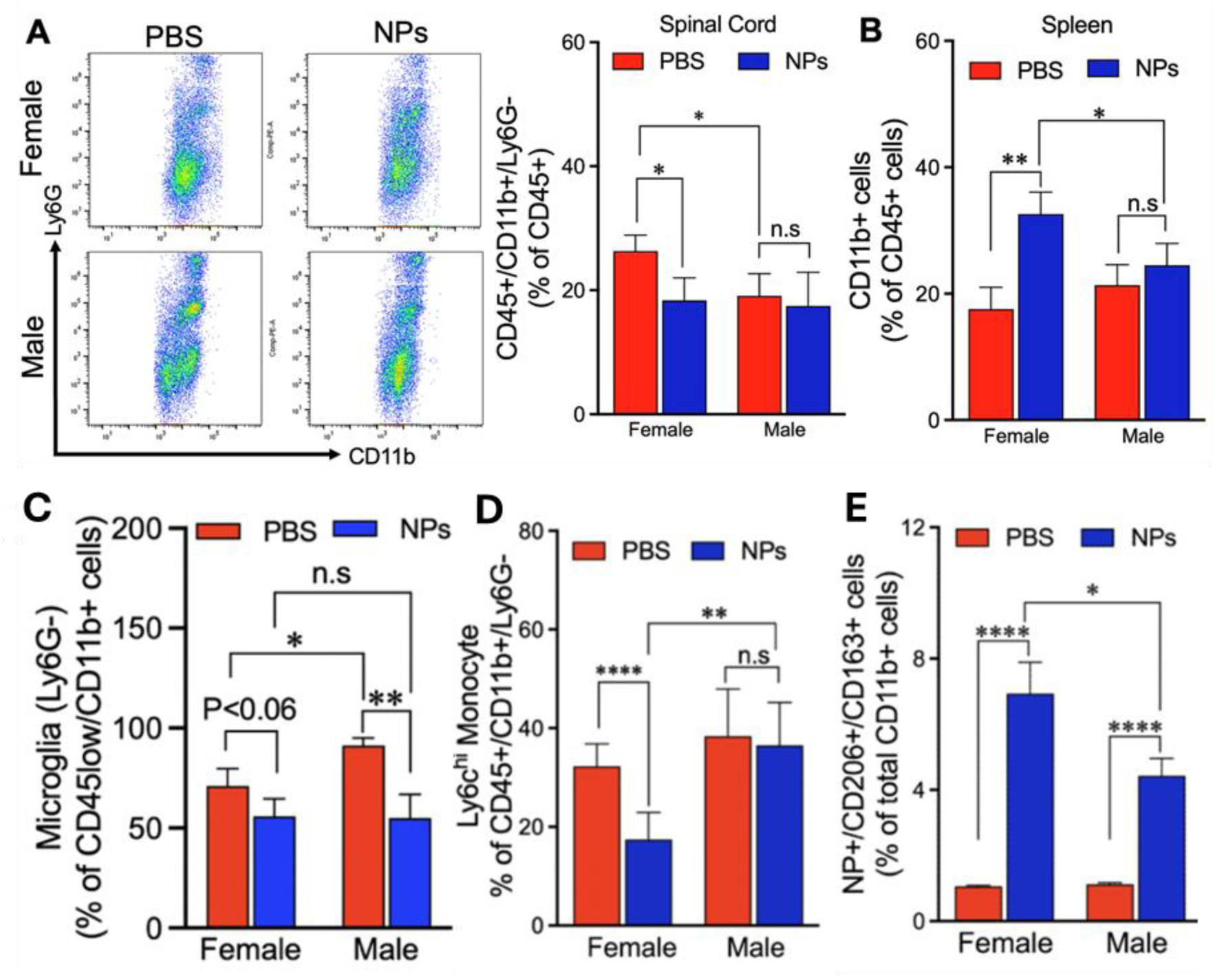
Sex-dependent effects of PLGA NPs on immune cell populations. The spinal cord and spleen were collected at 7-DPI for flow cytometry. (A) The proportion of infiltrated MDMs and microglia in the spinal cord. NP treatment significantly decreased the proportion in females but not in males. The female PBS group showed a higher proportion compared to the male PBS group. (B) The proportion of myeloid cells in the spleen. NP treatment increased the proportion of myeloid cells in females but not in males, with the female NP group showing a higher proportion than the male NP group. (C) The proportion of microglia in the spinal cord. It was higher in the male PBS group than in the female PBS group, and NP treatment reduced it in males but not in females. (D) The proportion of Ly6C^hi^ monocytes in the spinal cord. NP treatment significantly reduced the proportion of Ly6C^hi^ monocytes exclusively in females. (E) NP-treated females showed a higher proportion of NP-associated pro-regenerative type cells in the spinal cord than males. A two-way ANOVA with Tukey’s post hoc test for multiple comparisons. n = 4-5/group. n.s: no significant; * p<0.05, ** p<0.01, and **** p<0.0001. Data are presented as mean ± SEM.

We further analyzed the proportion of resident microglia population (CD45^low^/CD11b^+^/Ly6G^-^) in the injured spinal tissue and found that the microglia were more abundant in males than in females (Fig. 3C), consistent with previous studies [11]. NP treatment significantly reduced the microglia fraction in males (p<0.01) but showed a modest decrease in females (p<0.06). This decreased activation of microglia reinforces the importance of the indirect effects of NPs, which can be the result of a less inflammatory environment by impaired trafficking of pro-inflammatory immune cells into the injury [28]. These results suggest that PLGA NPs specifically affect myeloid cells infiltration in females and resident microglia in males, thereby reducing sex-specific immune differences after SCI.

Consistent with these findings, NP-treated females showed reduced Ly6C^hi^ monocytes at 7-DPI (Fig. 3D) and markedly decreased neutrophils (Ly6G^+^) infiltration at 1-DPI (Fig. S5). These results further support that NPs treatment more strongly limits circulating immune cells infiltration in females.

Finally, we observed sex-dependent effects of NPs treatment in macrophage polarization (Fig. 3E). NP-treated females demonstrated a significantly higher proportion of anti-inflammatory macrophage phenotypes (CD206^+^/CD163^+^) compared to males. This data indicates that systemic NP treatment promotes a more pro-regenerative myeloid phenotype in females. Collectively, by enhancing anti-inflammatory macrophage responses in females and attenuating microglial activation in males, PLGA NPs shifted both sexes toward a more balanced, pro-regenerative immune profile after SCI. These sex-specific, yet coordinated actions reduced the different inflammatory patterns typically observed between sexes and can account for the comparable functional recovery after NPs treatment across sexes.

### PLGA NPs modulate the polarization of macrophage and microglia at the lesion site

To validate the flow cytometry results at the lesion epicenter, we performed immunofluorescence staining on spinal cord sections at 7-DPI. F4/80 and CD206 antibodies were used to confirm macrophage polarization (Fig. 4A). The immunofluorescence results were consistent with the flow cytometry data. We confirmed the colocalization of NPs-Cy5.5 and F4/80^+^ cells at the lesion (Fig. S6). The density of F4/80+ cells was comparable across groups, with no significant change following NP treatment in either sex (Fig. S7A). However, NP administration markedly increased CD206 expression at the injury site (Fig. 4B and C), with both sexes showing elevated CD206^+^ cell numbers and females exhibiting a larger relative increase. Consequently, the proportion of F4/80^+^/CD206^+^ macrophages was significantly upregulated in NP-treated groups of both sexes (p<0.0001; Fig. S7B).

**Figure 4.**
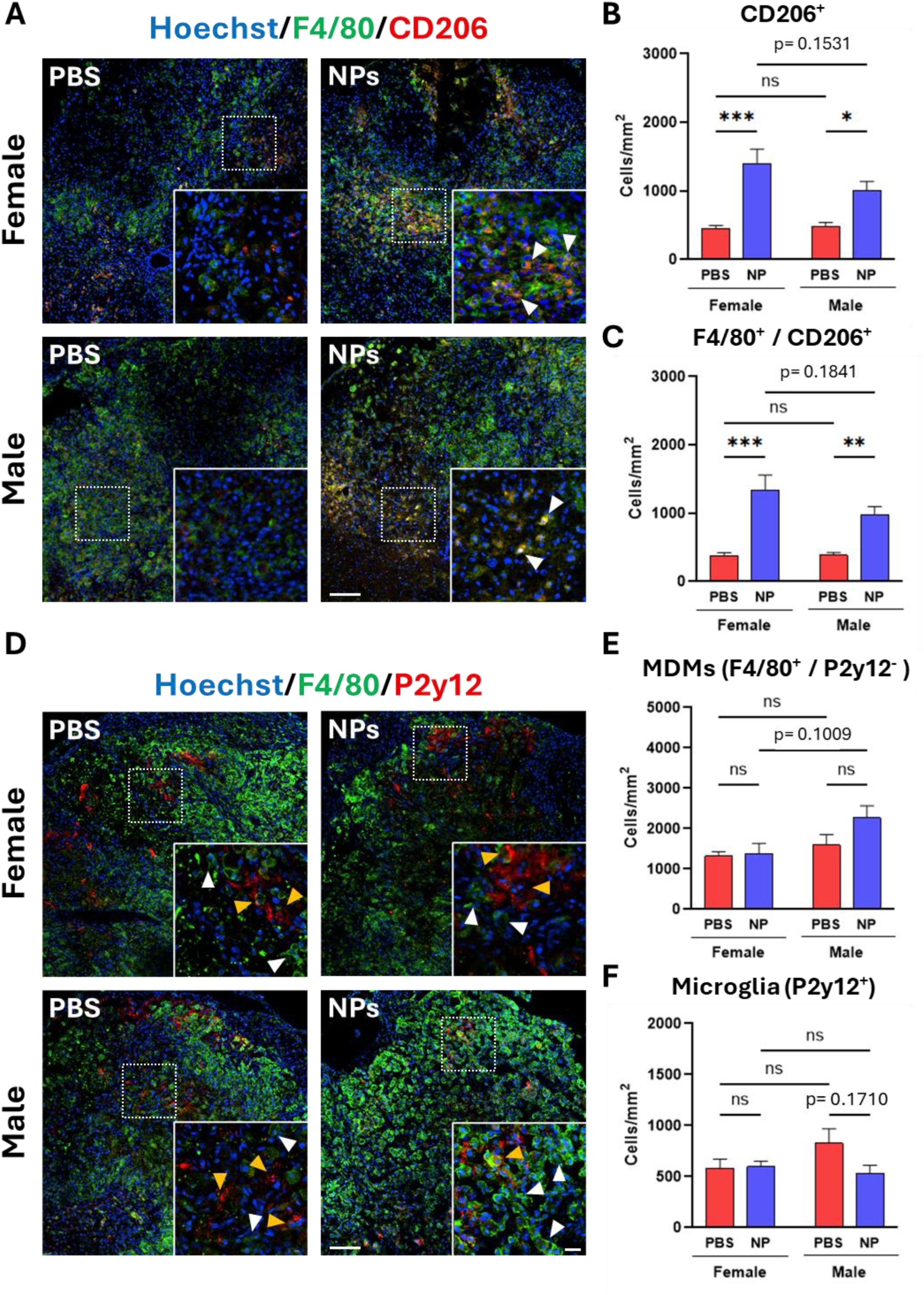
Immunofluorescence images for immune cells at 7-DPI. (A) Representative images of spinal cord sections were labeled using Hoechst, F4/80, and CD206. Scale bar = 100 µm. Boxed border regions were magnified, and anti-inflammatory (Hoechst^+^/F4/80^+^/CD206^+^) macrophages are represented with white arrowheads. Scale bar = 20 µm. (B) The density of total CD206^+^ cells. NP treatment upregulated the expression of CD206^+^ cells in both sexes. (C) The density of F4/80^+^/CD206^+^ cells. (D) Representative images of spinal cord sections were labeled using Hoechst, F4/80, and P2y12. Scale bar = 100 µm. Boxed border regions were magnified. MDM (F4/80^+^/P2y12^-^) and microglia (P2y12+) are represented with white and yellow arrowheads, respectively. Scale bar = 20 µm. (E) The density of MDMs (F4/80^+^/P2y12^-^). No changes were observed between groups. (F) The density of microglia (P2y12^+^). NP treatment reduced the density of microglia in males, but the change was not significant. A two-way ANOVA with Tukey’s post hoc test for multiple comparisons. n = 5/group. ns: no significant; * p<0.05, ** p<0.01, and *** p<0.001. Data are presented as mean ± SEM.

In contrast, immunofluorescence staining with F4/80 and P2y12 demonstrated sex-biased trends in MDM and microglia responses (Fig. 4D). NP-treated males tended to diminish the density of P2y12^+^ microglia and increase MDM density at the lesion site, whereas these populations were largely unaffected in females (Fig. 4E and F). Although there was no statistical significance, the observed trends align with flow cytometry findings, demonstrating that NP delivery preferentially suppresses resident microglia in males, while enhancing reparative macrophage polarization in females.

### PLGA NPs showed sex-specific regulation of inflammation-related gene expression

Having established sex-dependent immune responses to NP treatment, we next examined how these effects influence the transcriptional level. Total spinal cord RNA was analyzed utilizing the NanoString mouse inflammation panel and RT-qPCR.

Cell type profiling scores reflect the relative abundances of each cell population, determined by the expression of genes assigned to that cell type [29]. The female PBS group showed a slightly higher macrophage score compared to other groups, although the difference was not statistically significant (Fig. S8A). Notably, NP treatment reduced the macrophage score only in females, consistent with the flow cytometry findings. Moreover, the scores for dendritic cells, another phagocytic cell that is able to engulf PLGA NPs [30], showed a decreasing trend after NP treatment in both sexes (Fig. S8B).

Hierarchical clustering revealed distinct gene expression patterns depending on sex and NP treatment (Fig. 5A). Pathway profiling demonstrated pronounced sex differences at 7-DPI (Fig. 5B and S9). In PBS-treated mice, males showed broadly elevated inflammatory pathway activation compared with females, including chemokine receptor bind chemokines, interleukin-1 (IL-1) family signaling, and metabolism. NPs treatment markedly reduced these sex differences, indicating that males showed broad decreases across inflammatory signatures, whereas females showed only minimal transcriptional shifts.

**Figure 5.**
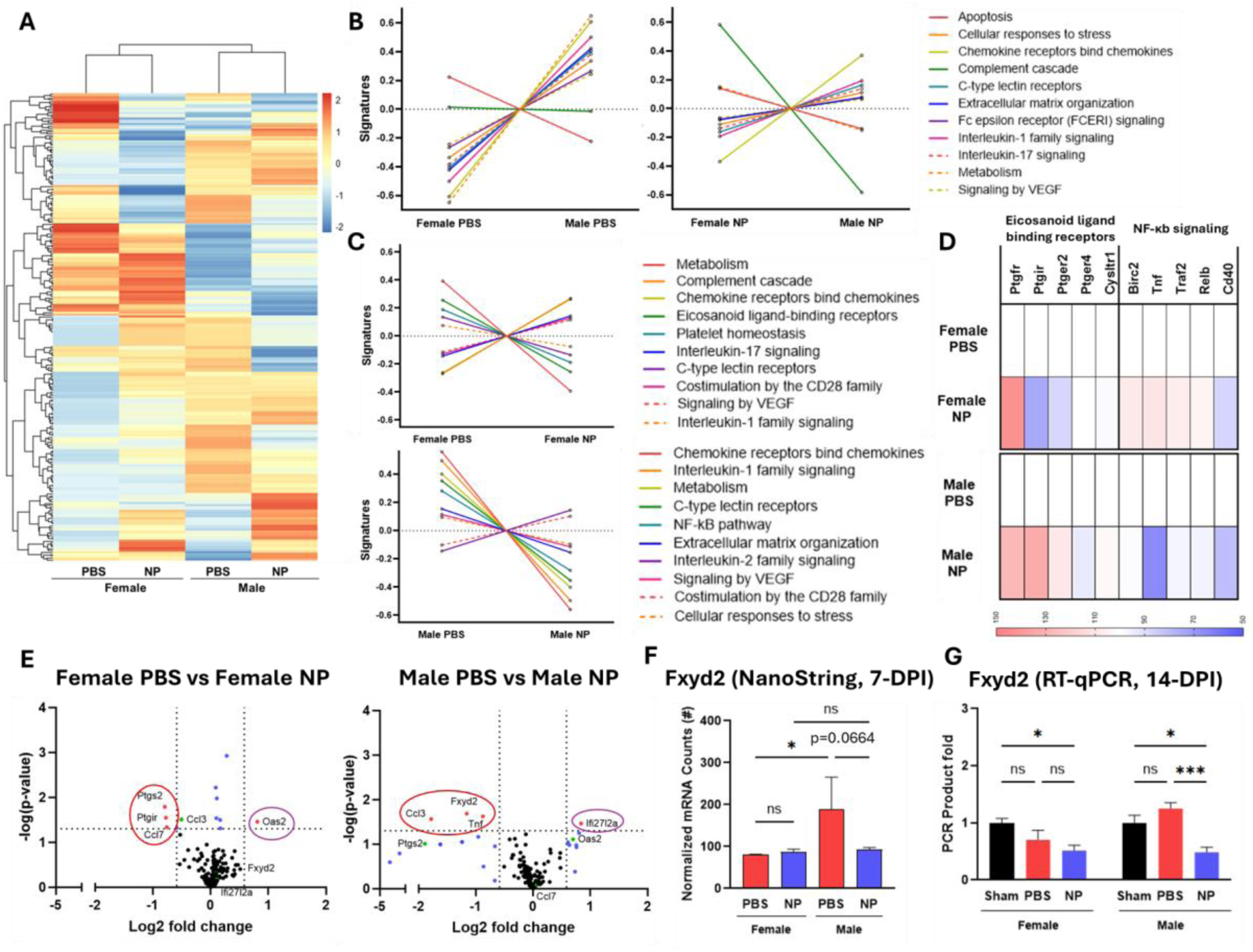
Gene expression analysis by NanoString and RT-qPCR. (A) Hierarchical clustering revealed distinct gene expression patterns depending on sex and NP treatment. (B) Pathway scores between males and females in PBS and NP-treated groups. NPs substantially reduced sex-dependent differences across most inflammatory pathways. (C) Pathway score plots display the top ten pathways most altered by NPs within each sex, highlighting distinct, sex-specific patterns of pathway modulation. (D) Heat maps of the genes in the pathways most responsive to NP treatment in males and females, revealing differential responses between sexes. (E) Volcano plots of changes in inflammatory gene expression of PBS versus NP-treated group in both sexes. Red: Genes with p-values less than 0.05 and > 1.5-fold change. Blue: Genes with p-values less than 0.05 or > 1.5-fold change. Green: Genes that are significantly changed in the other sex. n = 3/group. (F) Normalized mRNA counts of Fxyd2 from NanoString analysis. (G) Gene expression of Fxyd2 via RT-qPCR at 14-DPI. For NanoString, n = 3/group. For RT-qPCR, n = 3-4/group. A two-way ANOVA with Tukey’s post hoc test for multiple comparisons. ns: no significant; * p<0.05 and *** p<0.001. Data are presented as mean ± SEM.

Within-sex comparisons confirmed that NPs administration modulated distinct immune pathways in each sex (Fig. 5C). In females, NPs treatment markedly altered pathways related to metabolism, eicosanoid ligand-binding receptors, the complement cascade, and chemokine receptor bind chemokines. In males, pathways associated with chemokine receptor bind chemokines, IL-1 family signaling, metabolism, C-type lectin receptors, and NF-κB signaling were prominently affected by NPs. Consistent with these patterns, the directed global significance score, which reflects the overall magnitude and direction of gene expression changes within each pathway, identified the eicosanoid ligand-coupled receptor pathway as the most downregulated pathway in females (Score: -1.535) and the NF-κB signaling pathway as the most downregulated pathway in males (Score: - 1.719; Table. S3). Collectively, these findings suggest that NPs preferentially targets the eicosanoid signaling in females and NF-κB pathway in males.

Heatmaps of the pathway-associated genes demonstrated clear sex-dependent transcriptional alteration in responses to NP treatment (Fig. 5D). Moreover, we observed differentially expressed genes (DEGs) in volcano plots by four pairwise comparisons (Fig. 5E and S10). In females, NP treatment upregulated Oas2 and downregulated Ptgir, Ptgs2, and Ccl7, whereas in males, NPs upregulated Ifi27l2a and downregulated Tnf, Ccl3, and Fxyd2 (Fig. 5E).

Since Ptgir and Ptgs2 regulate eicosanoid ligand-coupled receptor pathway and eicosanoid synthesis [31] and Tnf, Ccl3, and Fxyd2 are associated with the NF-κB signaling [32, 33], these results are consistent with the pathway analysis, confirming sex-specific molecular responses to NP treatment. Collectively, these NanoString data indicate that PLGA NPs reprogram the lesion microenvironment via distinct transcriptional mechanisms in males and females.

Among the DEGs, Fxyd2 showed a significant sex-dependent difference at 7-DPI in both normalized mRNA counts and volcano plot comparing PBS-treated males and females (Fig. 5F and S10). Furthermore, RT-qPCR analysis at 14-DPI demonstrated that Fxyd2 expression remained significantly reduced in males following NP treatment, confirming the sustained impact of NPs on gene expression regulation (Fig. 5G and S11). Collectively, these results show that PLGA NPs act on innate immune cells at both transcriptional and cellular levels, reprogramming distinct but complementary inflammatory mechanisms in each sex.

### PLGA NPs enhanced axonal regrowth and remyelination, with sex-dependent differences in Schwann cell-mediated myelination

We investigated the impact of NP treatment on axonal regrowth and myelination in the chronic phase to confirm the functional recovery at the cellular level. Spinal cord sections at 56-DPI were stained using neurofilament 200 (NF200), myelin basic protein (MBP), and myelin protein zero (P0).

Immunofluorescence revealed improved tissue preservation and repair in NP-treated mice compared with PBS controls (Fig. 6A). NF200^+^ axons in NP-treated spinal cords were coated with MBP, suggesting productive remyelination. Quantitative analysis confirmed sex-dependent differences. Both sexes exhibited increased NF200 and MBP intensity after NP treatment, but the increase in MBP was significant only in females (Fig. 6B). Consistently, the number of NF200^+^ axons was significantly increased in NP-treated females (p<0.05), but not males (p=0.1029; Fig. 6C).

**Figure 6.**
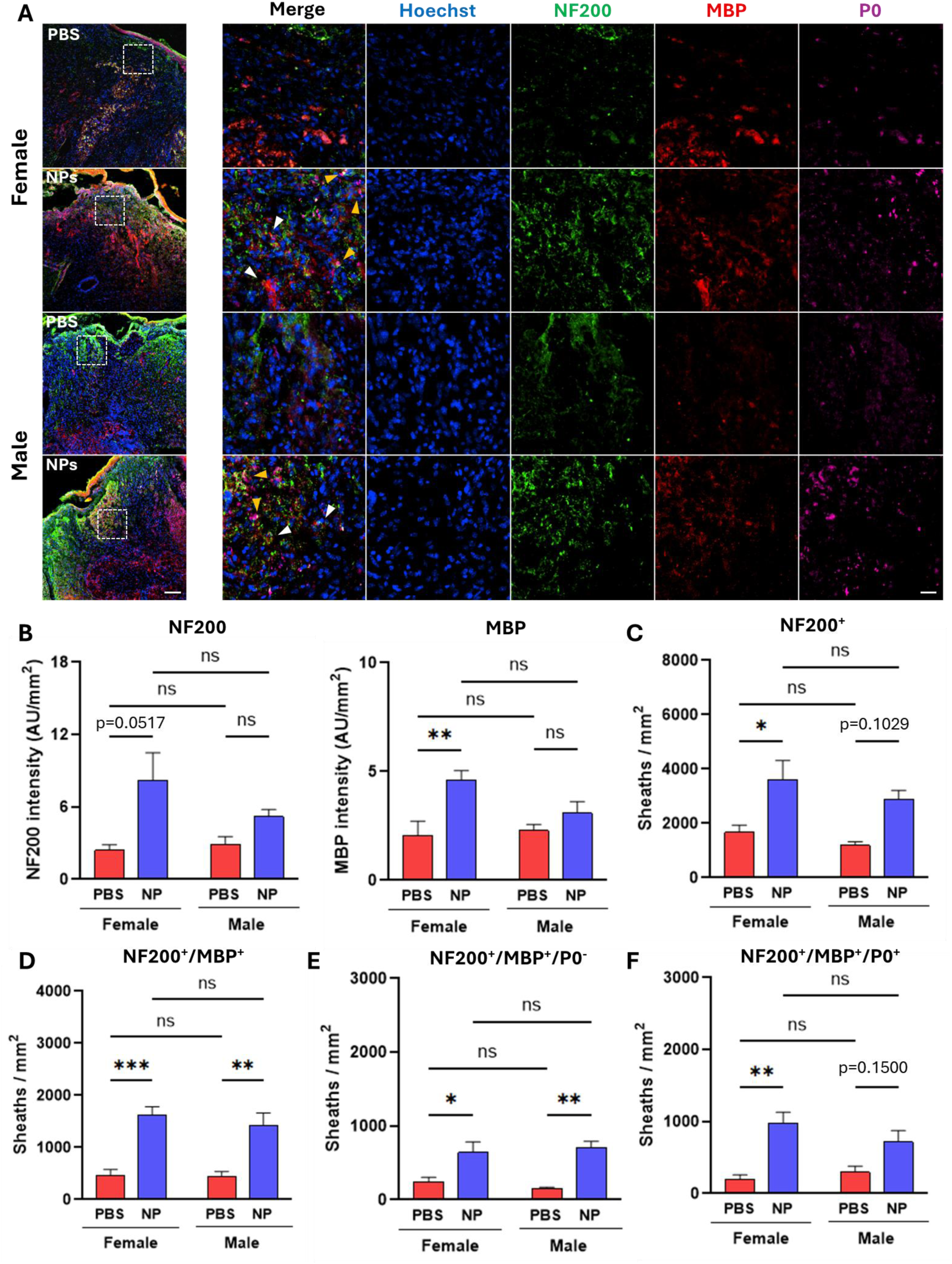
Axonal regrowth and remyelination in the chronic SCI phase. (A) Representative images of immunofluorescence staining. Spinal cord sections stained with Hoechst (blue), anti-NF200 (green), anti-MBP (red), and anti-P0 (magenta). Scale bar=100 µm. Boxed border regions were magnified, and oligodendrocyte-mediated myelinated axons (NF200^+^/MBP^+^/P0^−^), and Schwann cell-mediated myelinated axons (NF200^+^/MBP^+^/P0^+^) are represented with white and yellow arrowheads, respectively. Scale bar=20 µm. (B) The mean intensity of NF200 and MBP. (C to F) Quantification of the total number of axons (NF200^+^), myelinated axons (NF200^+^/MBP^+^), oligodendrocyte-mediated myelinated axons (NF200^+^/MBP^+^/P0^−^), and Schwann cell-mediated myelinated axons (NF200^+^/MBP^+^/P0^+^). NP treatment significantly increased Schwann cell-mediated myelinated axons only in females. N = 4-5/group. A two-way ANOVA with Tukey’s post hoc test for multiple comparisons. ns: no significant; * p<0.05, ** p<0.01, and *** p<0.001. Data are presented as mean ± SEM.

The number of myelinated axons (NF200^+^/MBP^+^) increased significantly after NP treatment in both sexes (Fig. 6D). To determine the source of myelination, P0 staining was used to differentiate between oligodendrocyte- and Schwann cell-derived myelin. Both female and male NP-treated groups showed a significant increase in oligodendrocyte-mediated myelinated axons (p<0.05 and p<0.01; Fig. 6E), indicating enhanced central nervous system remyelination. Interestingly, Schwann cell-mediated myelinated axons (NF200^+^/MBP^+^/P0^+^) were significantly increased only in females (p<0.01) but not in males (p=0.1500; Fig. 6F). Because Schwann cells infiltrate the lesion after SCI, regenerate peripheral myelin, and secrete neurotrophic factors that support axonal survival and growth [34-36], this sex-specific enhancement is consistent with a greater contribution of Schwann cell-associated repair processes in females. Overall, NP treatment promoted axonal regrowth and remyelination after SCI, with a preferential enhancement of Schwann cell-associated myelination in females.

### NP treatment decreases fibrotic scarring after SCI, but not in gliotic scarring

We investigated both fibrotic and gliotic scarring formation after NP treatment, because the scar tissue after SCI acts as a mechanical and chemical barrier to axon elongation [37, 38].

At 56-DPI, spinal cord sections were stained for fibronectin and glial fibrillary acidic protein (GFAP) to evaluate fibrotic and gliotic scars, respectively. In the sham group, we confirmed the absence of fibrotic and gliotic scarring (Fig. S12). In both sexes, NP treatment substantially reduced the area of fibronectin staining (Fig. 7A). The fibrotic scar core was smaller in NP-treated groups than in PBS groups, but the mean fibronectin intensity was significant only in female (p<0.01; Male: p=0.0503; Fig. 7B). In the analysis by sex, female mice tended to have a slightly lower fibronectin intensity than males in both PBS and NP-treated groups, although these were not statistically significant.

**Figure 7.**
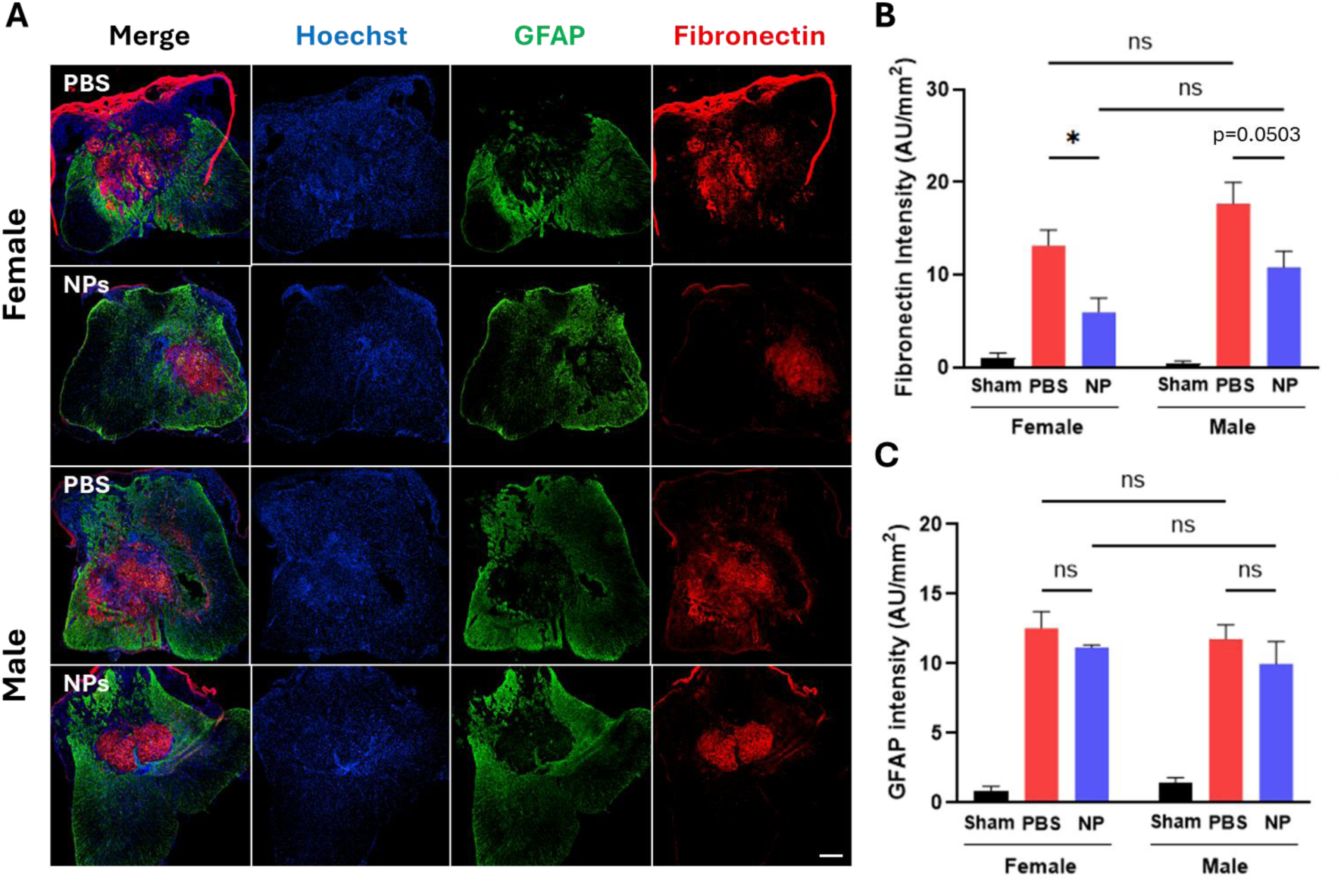
NP treatment reduces fibrotic but not gliotic scarring in chronic SCI. (A) Representative immunofluorescence images of spinal cord sections stained with anti-GFAP (green) and anti-fibronectin (red). NP treatment markedly reduced fibronectin-positive areas in both sexes. Scale bar = 200 µm. (B and C) Quantification of fibrotic (B) and gliotic (C) scarring. NP treatment substantially decreased fibrotic scarring, whereas gliotic scarring was not altered. N = 4-5/group. A two-way ANOVA with Tukey’s post hoc test for multiple comparisons. ns: no significant; * p<0.05. Data are presented as mean ± SEM.

In contrast, GFAP staining was comparable between NP-treated and PBS groups, and quantification confirmed no significant difference in mean GFAP intensity in either sex (Fig. 7A and C). Thus, PLGA NP treatment selectively attenuated fibrotic scar formation without altering reactive gliotic scarring, suggesting that the therapeutic effects of NPs primarily involve suppression of fibrosis rather than modulation of astroglial reactivity.

## Discussion

Our study demonstrated that systemic administration of PLGA NPs after SCI enhances recovery in both sexes through sex-specific immune reprogramming. NP treatment significantly improved locomotor outcomes in both sexes. Notably, significant sex differences were observed in PBS-treated controls, consistent with previous reports of sexually dimorphic recovery after SCI [39, 40]. In PBS groups, females achieved higher BMS sub-scores at later time points than males, reflecting an intrinsic advantage in functional recovery after SCI. Importantly, NP treatment eliminated this disparity, with NP-treated males achieving locomotor scores comparable to those of NP-treated females. These findings indicate that PLGA-NPs not only promote overall functional recovery but also mitigate baseline sex differences in regenerative outcomes after SCI.

This convergence in outcomes results from sex-specific interaction between NPs and the immune system. Although PLGA NPs robustly modulate the post-SCI inflammatory environment in both sexes, the nature of this modulation differs between males and females. Female exhibited greater NP accumulation in the spleen and reduced NP presence within the injured spinal cord compared with males. Consistent with this distribution, NPs treatment significantly increased CD11b^+^ cells in females and corresponding reduction in Ly6C^hi^ monocytes infiltration at the lesion site. In contrast, males exhibited lower splenic NP accumulation, greater NP presence within the injured spinal cord, and a reduction in resident microglial density.

These sex-specific patterns of NPs biodistribution and immune cell modulation suggest that post-injury inflammation in females is driven primarily by infiltrating MDMs, whereas males rely more heavily on microglial activation. This distinction is consistent with previous reports showing enhanced recruitment of MDM in females and greater dependence on resident microglia in males after SCI [10, 11]. By selectively engaging these dominant immune populations, NP treatment rebalances sex-biased innate immune responses and reshapes the inflammatory landscape after injury. Furthermore, consistent with this immune reprogramming, both immunofluorescence and flow cytometry data demonstrated an increase in CD206^+^ cell populations after NP treatment, reflecting a shift toward a pro-regenerative phenotype. This increase was more pronounced in females, in line with prior evidence that female microglia exhibit higher CD206 expression than males [11]. Collectively, these findings suggest that female myeloid cells may have a greater intrinsic capacity for regenerative phenotype after SCI, which is further amplified by NPs-mediated immunomodulation.

While flow cytometry and immunofluorescence analyses showed broadly consistent patterns, minor discrepancies were observed in infiltrated immune cells quantification. Flow cytometry revealed a reduction in infiltrating macrophages in NP-treated females, whereas immunofluorescence analysis did not show a corresponding decrease in F4/80^+^ cells at the lesion epicenter. This divergence likely reflects the different experimental methods. Flow cytometry analysis uses the spinal cord tissue encompassing multiple levels, thereby capturing both lesion core and perilesional responses, whereas immunofluorescence provides a more spatially restricted assessment of the epicenter. The use of distinct cellular markers may further contribute to these discrepancies. In addition, NP treatment may preferentially reduce inflammatory cell accumulation in surrounding tissue by limiting lesion expansion, without substantially altering the dense macrophage population at the lesion epicenter.

To further link these cellular effects to underlying molecular mechanisms, we performed NanoString analysis to assess transcriptional modulation by NPs treatment. Notably, inflammatory pathways that exhibited sex-dependent differential expression after SCI showed reduced divergence following NP administration. These findings indicate that post-injury inflammatory gene expression differs between males and females, and that NP treatment mitigates these transcriptional differences. This transcriptional convergence is consistent with the observed functional outcomes, in which NP treatment diminished baseline sex-based differences in recovery.

Moreover, NanoString data further revealed that PLGA NPs induce distinct, sex-specific patterns of inflammatory gene regulation. In females, NP treatment downregulated Ptgir and Ptgs2, indicating suppression of cyclooxygenase-mediated eicosanoid signaling, a pathway that drives prostaglandin E₂ (PGE₂) production and amplifies inflammation after injury [41]. This downregulation indicates a shift toward a less inflammatory transcriptional state in females. In contrast, NP treatment in males primarily downregulated expression of Tnf and Ccl3, two key pro-inflammatory cytokines downstream of NF-κB signaling [32].

Collectively, these transcriptional changes closely align with the immune population profiles observed by flow cytometry. The eicosanoid signaling regulates macrophage and neutrophil activation and chemotaxis [42, 43], while NF-κB signaling pathway controls pro-inflammatory cytokine production and microglia activation [44, 45]. In females, downregulation of eicosanoid- and chemokine-related genes (Ptgir, Ptgs2, Ccl7) correlated with reduced MDM and neutrophil infiltration, alongside enhanced polarization into pro-regenerative type. In males, suppression of NF-κB–associated genes (Tnf, Ccl3) coincides with diminished microglial activation, consistent with the known role of activated microglia as major producers of Tnf and Ccl3 [46]. This suppression of NF-κB-associated genes is further supported by the known immunomodulatory effects of lactic acid, a degradation byproduct of PLGA NPs, which inhibits NF-κB activation and nuclear translocation [47]. Together, these findings indicate that PLGA NP treatment modulates innate immune responses at both molecular and cellular levels through sex-dependent yet convergent mechanisms, ultimately fostering a pro-regenerative milieu and minimizing sex-specific inflammatory disparities after SCI.

Furthermore, our gene expression analysis identified Fxyd2 as an additional sex-specific target of NP treatment. At 7-DPI, Fxyd2 expression was significantly higher in the male PBS group compared to the female PBS group. NP treatment selectively downregulated Fxyd2 in males, and this reduction persisted up to 14-DPI, whereas no significant change was observed in females. Fxyd2, associated with the NF-κB signaling pathway [33], encodes the γ-subunit of the Na⁺/K⁺-ATPase (NKA) and known to be upregulated under conditions of cellular stress and inflammation [33, 48]. Therefore, elevated Fxyd2 expression in males may reflect upregulated inflammatory activation associated with increased NF-κB signaling in microglia. Consistent with this, our data demonstrate that NP treatment reduced both microglia activation and Fxyd2 expression specifically in males. This coordinated reduction indicates that NP-mediated inhibition of the NF-κB pathway in microglia attenuates inflammatory activation that normally induces Fxyd2 upregulation. In here, Fxyd2 likely functions as a downstream transcriptional indicator of NF-κB-associated microglial activation rather than as a direct regulator of inflammatory signaling.

Beyond its function in inflammation, the regulation of Fxyd2 also had important implications for neuronal hyperexcitability and pain sensitization. Previous studies have shown that loss of Fxyd2 attenuates mechanical hypersensitivity in neuropathic pain models [49], and Fxyd2-deficient mice exhibit an accelerated resolution of inflammatory allodynia compared to wild-type controls [50]. Therefore, the sex-specific expression of Fxyd2 and its selective modulation by NP treatment highlight the need for further investigation into how Fxyd2 contributes to post-SCI inflammatory signaling and neural excitability, and whether NP-medicated immune reprogramming offers sex-dependent benefits in neuropathic pain following SCI.

The molecular and immune reprogramming induced by NP treatment during the acute phase was associated with improved anatomical and functional restoration during the chronic phase. NP treatment promoted neural regeneration at the lesion site, with females demonstrating greater axonal regeneration compared to males. Notably, females also showed a significant increase in peripheral (Schwann cell-mediated) remyelination relative to males. These effects may be associated with sex-specific differences in pro-regenerative macrophage responses. Pro-regenerative macrophages secret anti-inflammatory cytokines and trophic factors such as IL-10, TGF-β, NGF, and BDNF, which promote Schwann cell proliferation and establish a permissive microenvironment for repair [51]. Accordingly, the greater expansion of pro-regenerative macrophage populations observed in females following NP treatment may contribute to the enhanced Schwann cell–mediated remyelination in this group. In contrast, oligodendrocyte-mediated remyelination after NP treatment appeared largely sex-independent, potentially reflecting temporal differences between remyelination pathways. Oligodendrocyte-mediated remyelination typically begins more than two weeks post-injury, whereas Schwann cell-mediated myelination occurs within the first two weeks and progresses at a relatively steady rate [52]. Thus, NP administration during the first 7 days after SCI and the resulting pro-regenerative immune microenvironment may preferentially influence Schwann cell-mediated remyelination. However, additional biological factors, including sex hormones and intrinsic cellular differences, may also contribute to remyelination dynamics and warrant further investigation.

Consistent with the observed enhancements in tissue repair, NP-treated mice exhibited a significant reduction in fibrotic scar formation, whereas gliotic scarring remained largely unaffected. This finding differs from our previous study, which demonstrated significant reduction in both fibrotic and gliotic scarring following PLGA NP treatment [17]. The discrepancy likely reflects differences in injury models and therapeutic approaches. In our prior work, PLGA NPs were delivered alongside a PLGA multichannel bridge in a hemisection injury model, whereas the present study employed a contusion model with systemic NP administration alone. Notably, the selective attenuation of fibrotic scarring observed here is consistent with previous reports indicating that PLGA NPs preferentially modulate fibrotic, but not astroglial scar formation [28]. In line with prior reports, infiltrating blood-derived monocytes and macrophages have been identified as key drivers of fibrotic scar formation after SCI. These circulating immune cells recruit perivascular fibroblasts into the lesion site, and depletion of hematogenous macrophages has been shown to reduce fibroblast accumulation and fibrotic scar area, thereby facilitating axonal regrowth [53]. Thus, the reduction in fibrotic scarring observed in our study likely reflects NP-mediated modulation of infiltrating immune cells without disrupting the astroglia barrier. Notably, the reduction in fibronectin intensity was more pronounced in the female group, suggesting stronger NP-mediated effects on peripheral myeloid populations. In females, greater NP accumulation in the spleen was associated with reduced recruitment of pro-inflammatory and pro-fibrotic macrophages to the injured spinal cord, alongside downregulation of eicosanoid- and chemokine-related gene expression. These alterations are expected to limit macrophage-fibroblast interactions that promote fibrotic scar formation. In contract, in males, NPs dominantly targeted resident microglia, suppressed NF-κB associated signaling pathway without altering in peripheral macrophage infiltration. Given that fibrotic scarring is driven predominantly by infiltrating macrophages rather than microglia, this sex-specific pattern of immune modulation provides a plausible explanation for the greater reduction in fibrotic scarring observed in females compared with males.

## Conclusions

In summary, our study demonstrates that systemic administration of PLGA NPs after SCI promotes recovery through sex-dependent immune reprogramming. NP treatment reshaped inflammatory cell responses in distinct ways in males and females, leading to reduced scar formation, enhanced axonal regeneration and remyelination, and improved functional outcomes. Although the underlying cellular and molecular mechanisms differed between sexes, reflecting differential engagement of peripheral macrophage- and microglia-driven inflammatory pathways, the therapeutic outcomes converged across sexes. These findings establish drug-free polymeric NPs as an effective immunomodulatory strategy for SCI and underscore the importance of incorporating sex as a fundamental biological variable. Collectively, consideration of sex-dependent immune mechanisms will be critical for optimizing future nanotherapeutic approaches, with implications extending beyond SCI to other inflammation-mediated diseases.

## Acknowledgments

This work was supported by National Institutes of Health through R01NS136272, the National Center for Advancing Translational Sciences UL1 TR001998, the University of Kentucky Neuroscience Research Priority Area (NRPA017) and the Center for Pharmaceutical Research and Innovation NIH P20 GM130456. We would like to thank the University of Kentucky’s Flow Cytometry and Immune Monitoring Core Facility and Genomics Core Laboratory. The access of BioRender is supported by the Bioelectronics and Nanomedicine Research Center (BNRC). Graphic abstract was created in BioRender. Nano, B. (2026) https://BioRender.com/femldnf.

## Autor Contributions

**Jaechang Kim**: Conceptualization, Data curation, Writing – review & editing, Writing – original draft, Methodology, Investigation, Visualization. **Irina Kalashnikova**: Methodology, Investigation, Visualization. **Ruby Maharjan**: Methodology, Investigation, Visualization. **Fernanda Stapenhorst França**: Investigation. **Daniel Kolpek**: Investigation. **James Ogidi**: Investigation. **John C. Gensel**: Conceptualization, Writing—review & editing. **Jonghyuck Park**: Conceptualization, Funding acquisition, Project administration, Writing—review & editing, Supervision.

## Notes

### Competing Interest Statement

The authors have declared no competing interest.

