## Supplementary Material for "Systemic delivery of drug-free polymeric nanoparticles reprograms innate immunity in a sex-dependent manner after spinal cord injury"

Supplementary Figures

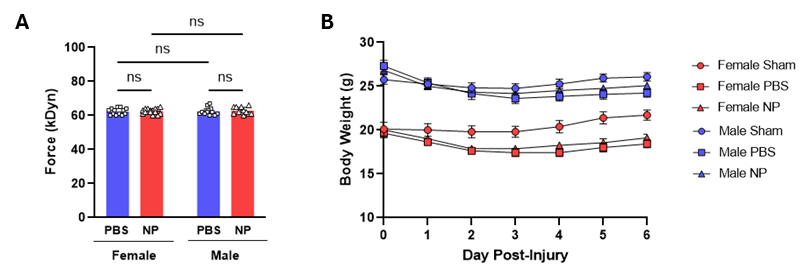

Fig. S1. Contusion device impact force and body weight change after SCI. (A) Contusion device impact force confirms reproducible and consistent injuries between groups. (B) Changes in body weight after SCI. No significant differences were observed between PBS and NP-treated groups. Male sham: n = 12; female sham: n = 10; male PBS: n = 13; female PBS: n = 14; male NP: n = 12; female NP: n = 16. A two-way ANOVA with Tukey’s post hoc test for multiple comparisons. ns; no significant. Data are presented as mean ± SEM.

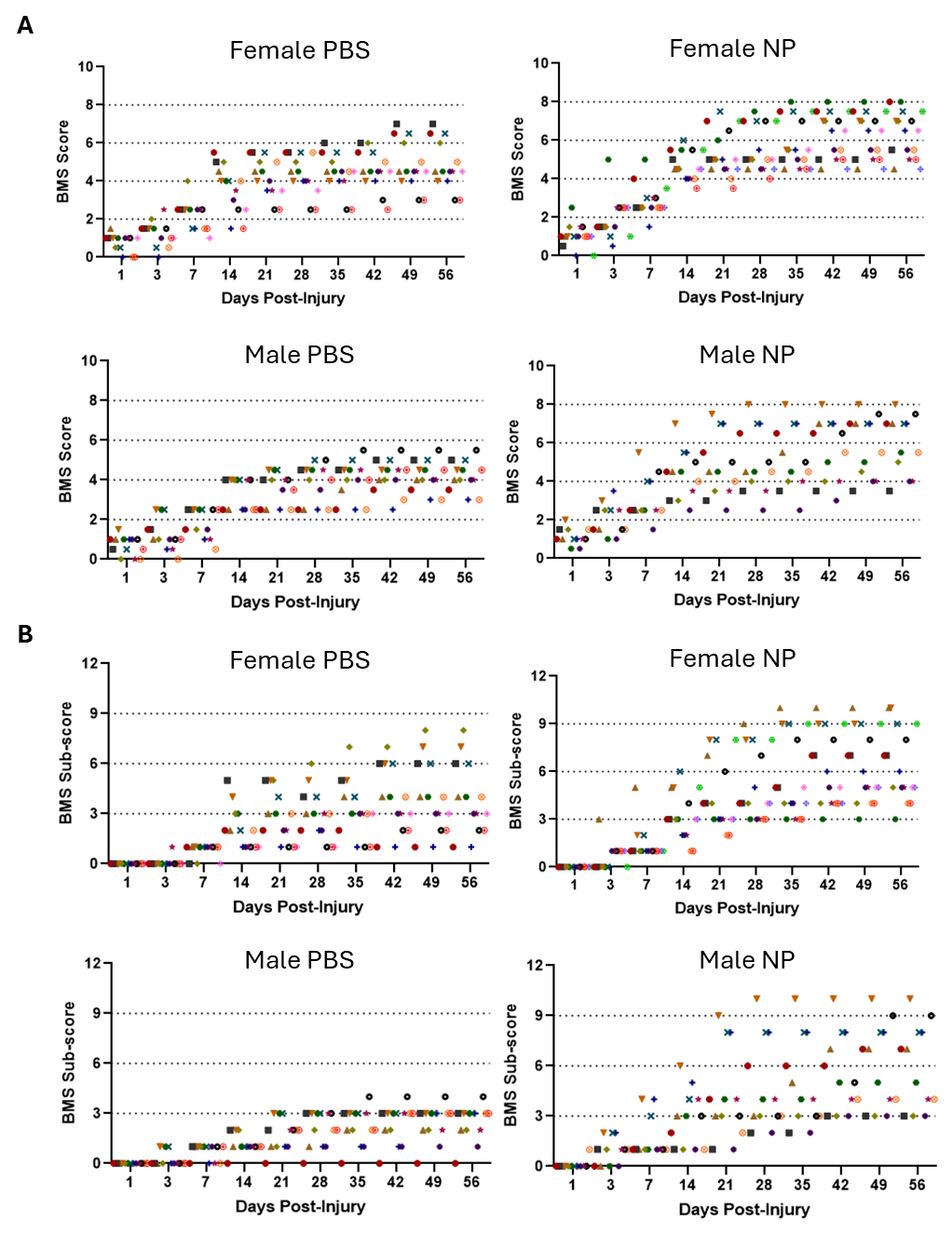

Fig. S2. Detailed BMS score and sub-score data. Scatter plot diagrams display (A) BMS score and (B) sub-score distributions of PBS and NP-treated groups in both sexes.

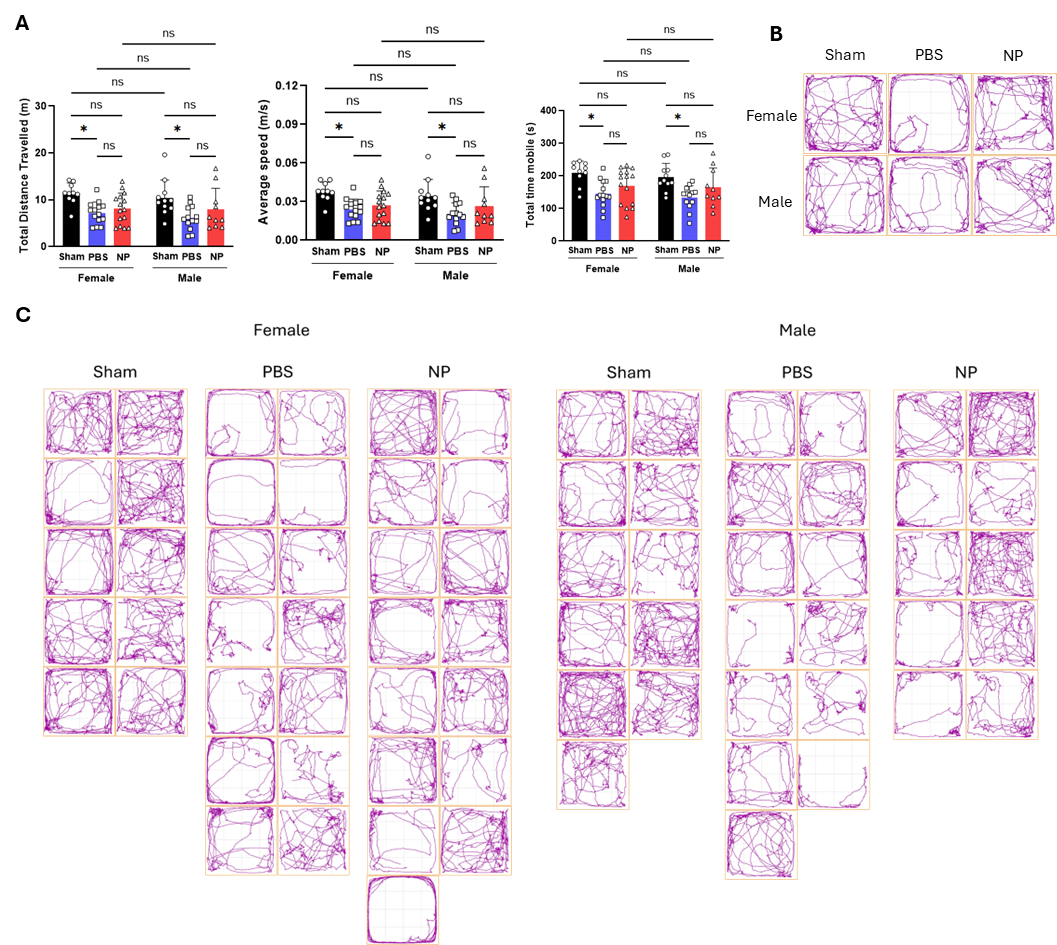

Fig. S3. Spontaneous activity in the open field test at 56-DPI. (A) Total distance traveled, average speed, and total time mobile at 56-DPI. An increasing trend was observed in NP-treated groups compared to PBS groups in both sexes. (B) Representative and (C) all images displaying traces of mice in the open field test, illustrating improved activity in NP-treated groups compared to PBS groups. Male sham: n = 11; female sham: n = 10; male PBS: n = 13; female PBS: n = 14; male NP: n = 10; female NP: n = 15. Two-way ANOVA with Tukey’s post hoc test for the multiple comparisons. ns: no significant; * p < 0.05. Data are presented as mean ± SEM.

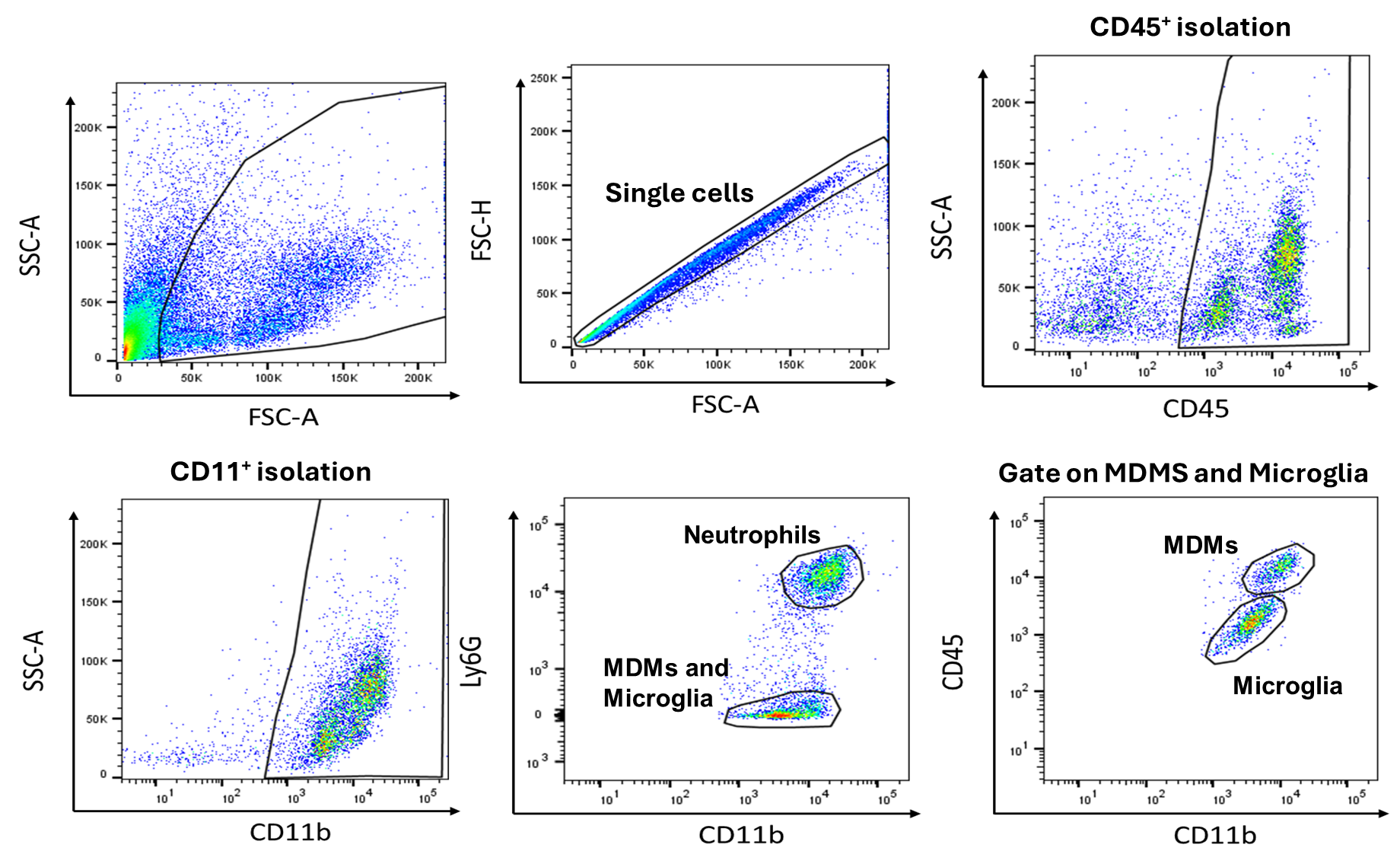

Fig. S4. Gating strategy used to identify the myeloid subpopulations. Cells were first gated according to FSC-SSC, then restricted to singles cells. Neutrophils were identified as CD45^+^/CD11b^+^/Ly6G^+^. The MDMs and microglia (CD45^+^/CD11b^+^/Ly6G^-^) were divided into MDMs (CD45^high^/CD11b^+^/Ly6G^-^) and Microglia (CD45^low^/CD11b^+^/Ly6G^-^).

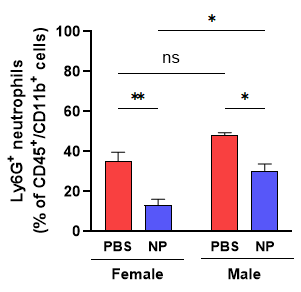

Fig. S5. PLGA NPs reduced neutrophil infiltration in the spinal cord at 1-DPI. The spinal cord was collected at 1-DPI for flow cytometry. The proportion of infiltrated neutrophils (Ly6G^+^) was significantly reduced by NP treatment, with a more pronounced effect observed in female mice. A two-way ANOVA with Tukey’s post hoc test for multiple comparisons. n = 4-5/group. n.s: no significant; * p<0.05 and ** p<0.01. Data are presented as mean ± SEM.

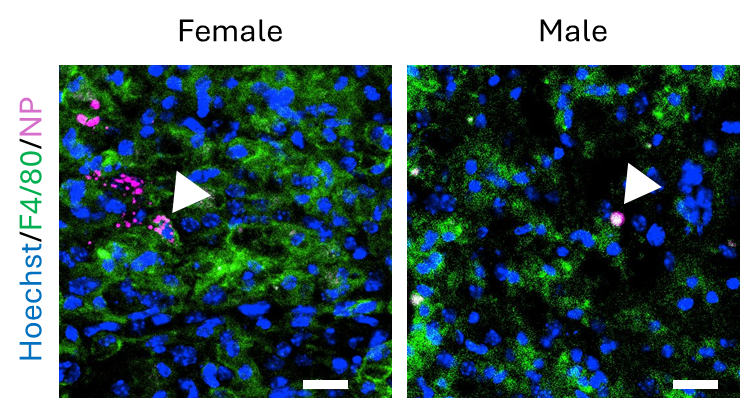

Fig. S6. NP colocalization and internalization with F4/80^+^ cells. Spinal cord sections at 7-DPI were labelled for Hoechst (blue), anti-F4/80 (green), and NPs-Cy5.5 (violet). NPs-Cy5.5 were colocalized with F4/80^+^ cells (white arrows). Scale bar = 20 µm.

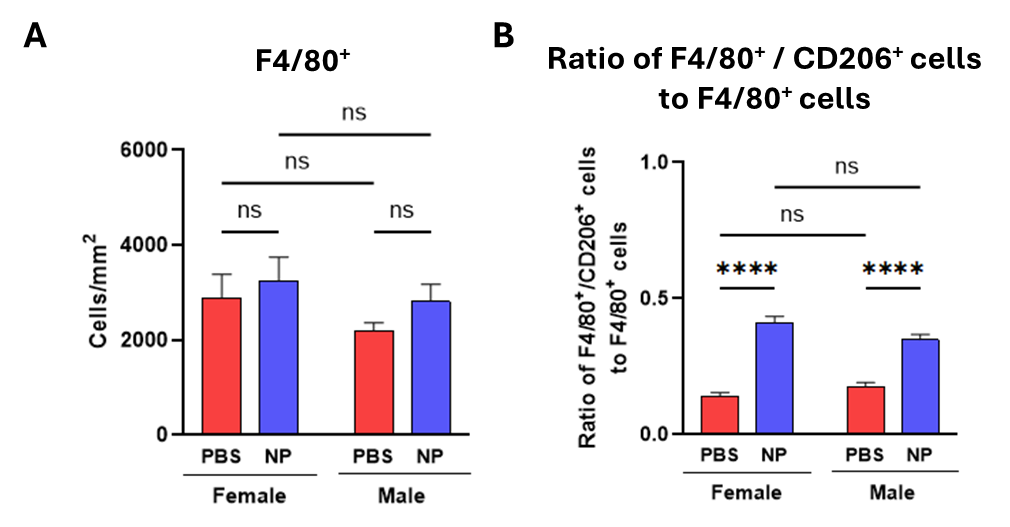

Fig. S7. Quantification of immunofluorescence images for immune cells at 7-DPI. (A) The density of total F4/80^+^ macrophages. No changes were observed between groups. (B) The ratio of F4/80^+^/CD206^+^ cells to total F4/80^+^ macrophages. A two-way ANOVA with Tukey’s post hoc test for multiple comparisons. n = 5/group. ns: no significant; * p<0.05, ** p<0.01, *** p<0.001, and **** p<0.0001. Data are presented as mean ± SEM.

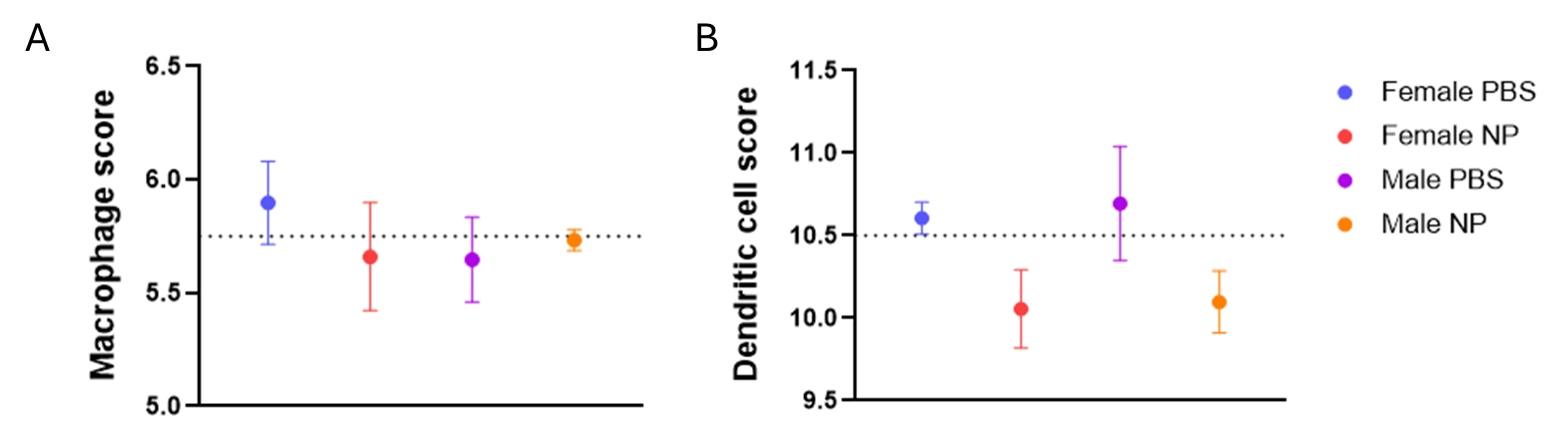

Fig. S8. Cell type profiling analysis generated using the Advanced Analysis module in nSolver. (A) Macrophage scores across groups. The female PBS group displayed a slightly higher macrophage score relative to other groups. (B) Dendritic cell scores, showing a decreasing trend after NP treatment in both males and females.

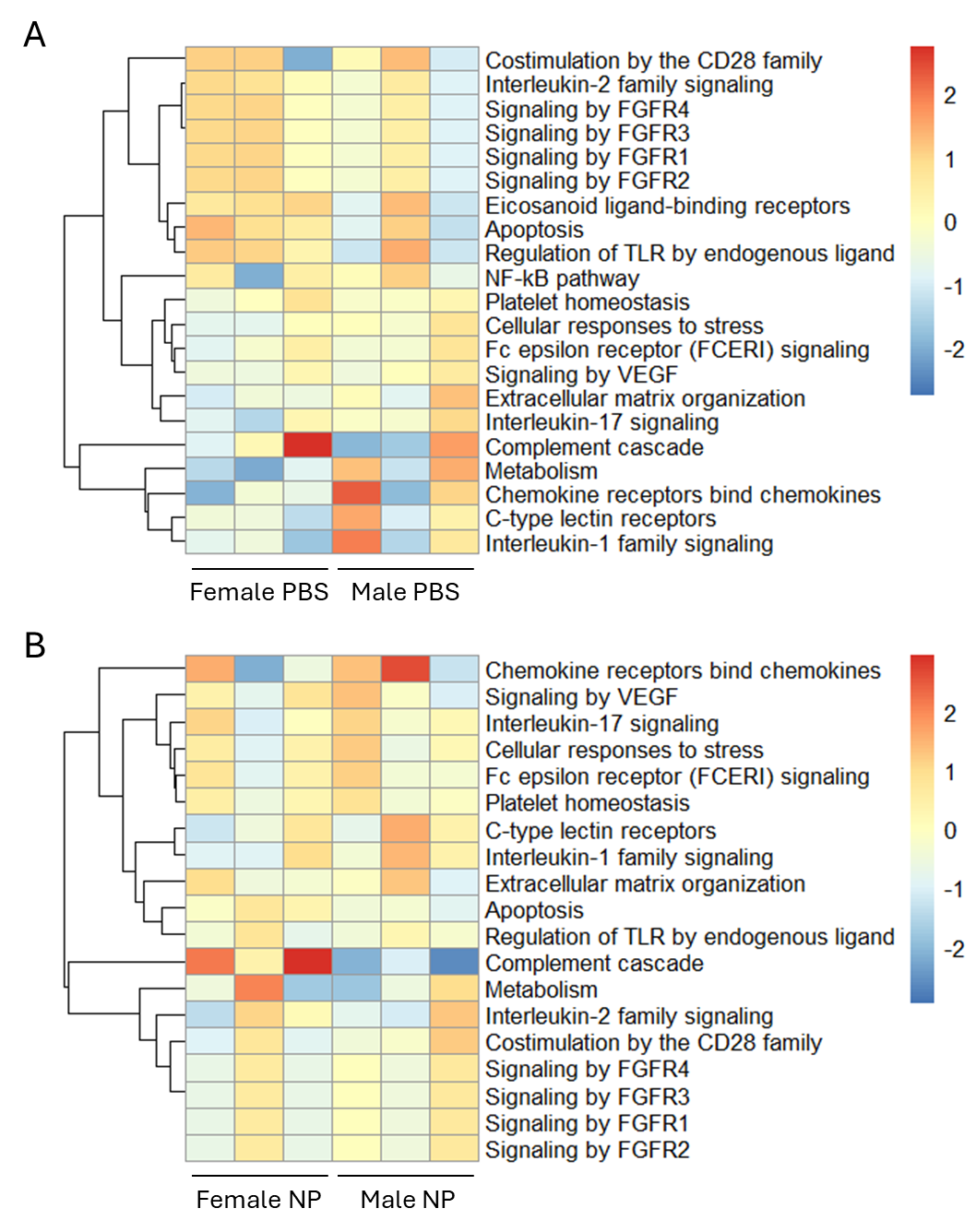

Fig. S9. Heatmap of pathway scores comparing males and females in PBS (A) and NP-treated (B) groups. Heatmap showing pathway scores to provide an overview of how pathway activity patterns vary across samples. Red indicates higher pathway scores, and blue indicates lower scores.

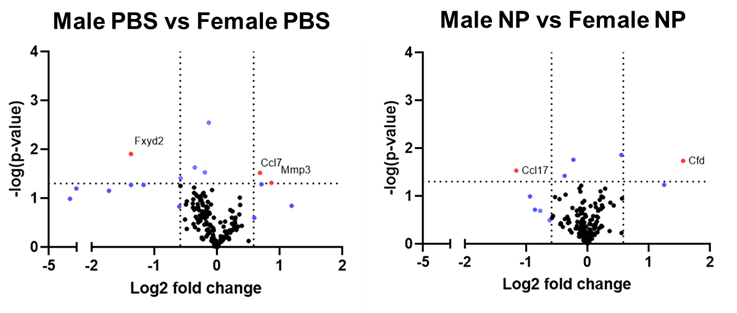

Fig. S10. Volcano plots of changes in inflammatory gene expression of males versus females. Volcano plots showed the different regulations of inflammatory genes between males and females. Red: Genes with p-values less than 0.05 and > 1.5-fold change. Blue: Genes with p-values less than 0.05 or > 1.5-fold change. N = 3/group.

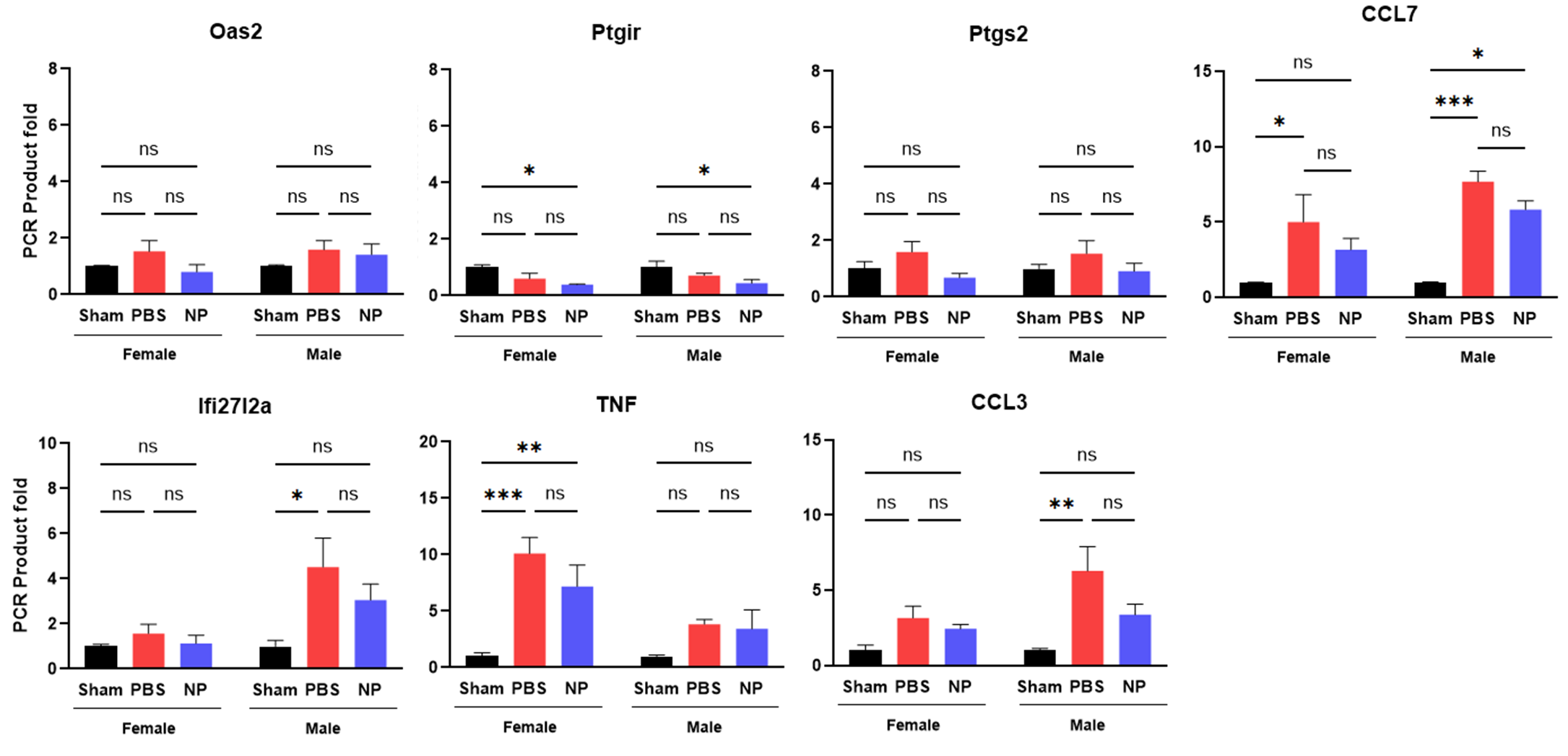

Fig. S11. Gene expression analysis by RT-qPCR at 14-DPI. The 14-DPI expression of differentially expressed genes (DEGs) identified in 7-DPI volcano plots between PBS- and NP-treated groups was confirmed by RT-qPCR in both sexes. A two-way ANOVA with Tukey’s post hoc test for multiple comparisons. n = 3-4/group. ns: no significant; * p<0.05, ** p<0.01, and *** p<0.001. Data are presented as mean ± SEM.

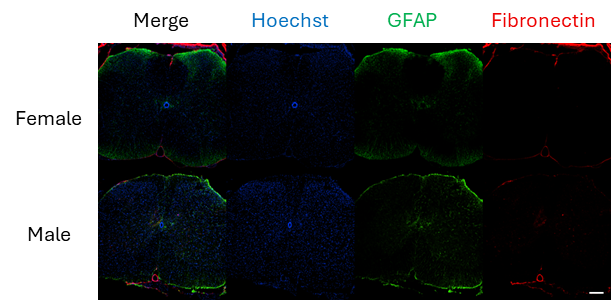

Fig. S12. Immunofluorescence image of sham groups at 56-DPI. Spinal cord sections were stained with Hoechst (blue), anti-GFAP (green), and anti-fibronectin (red). No fibrotic or gliotic scarring was observed in sham groups. Scale bar = 200 µm.

Supplementary Tables

Table S1. Primary and secondary antibodies for immunofluorescence

| **Name** | **Company** | **Catalog number** | **Concentration** |
| --- | --- | --- | --- |
| F4/80 | Abcam | ab6640 | 1:200 |
| CD206 | Abcam | ab64693 | 1:1000 |
| P2y12 | Abcam | ab300140 | 1:500 |
| Neurofilament 200 | Sigma | N4142 | 1:100 |
| Myelin basic protein | Abcam | Ab11159 | 1:500 |
| P0 myelin protein | Abcam | ab39375 | 1:100 |
| GFAP | Invitrogen | 13-0300 | 1:1000 |
| Fibronectin | Sigma | F7387 | 1:1000 |
| Hoechst 33342 | Thermo Fisher | 62249 | 1:2000 |
| Alex Fluor^TM^ 488 | Thermo Fisher | A21208 | 1:1000 |
| CF^TM^ 488, 555, and 633 | Sigma Aldrich | SAB4600061, SAB4600036, SAB4600060, and SAB4600127 | 1:1000 |

Table S2. Primer sequence for RT-qPCR

| **Gene** | **Accession number** | **Forward (5’-3’)** | **Reverse (5’-3’)** |
| --- | --- | --- | --- |
| Oas2 | NM_145227 | CACCAAAGTCCTGAAGACCGTC | AGAGTCGTAACTCTCCAGCGAG |
| Ptgir | NM_008967 | GTTTACCACCTGATTCTGCTGGC | CGTTGAAGCGGAAGGCGAGGA |
| Ptgs2 | NM_011198 | GCGACATACTCAAGCAGGAGCA | AGTGGTAACCGCTCAGGTGTTG |
| CCL7 | NM_013654 | CAGAAGGATCACCAGTAGTCGG | ATAGCCTCCTCGACCCACTTCT |
| Ifi27I2a | NM_029803 | CTTCACTGGGACAGGCATTGCA | CCTGCTGATTGGAGTGTGGCTA |
| TNF | U68416.1 | TTGACCTCAGCGCTGAGTTG | CCTGTAGCCCACGTCGTAG |
| CCL3 | NM_011337 | ACTGCCTGCTGCTTCTCCTACA | ATGACACCTGGCTGGGAGCAAA |
| Fxyd2 | NM_052823.2 | ACTATGAAACCGTCCGCAAA | TTCTTACCGCCCCCACAG |
| 18s-rRNA | NR_003278.3 | GCAATTATTCCCCATGAACG | GGCCTCACTAAACCATCCAA |

Table S3. Global significance Scores from NanoString

| **Male PBS vs Male NP** | **Undirected global significance score** | **Directed global significance score** |
| --- | --- | --- |
| NF-κB pathway | 1.719 | -1.719 |
| Signaling by FGFR1 | 1.374 | -1.314 |
| Signaling by FGFR2 | 1.374 | -1.314 |
| Signaling by FGFR3 | 1.374 | -1.314 |
| Signaling by FGFR4 | 1.374 | -1.314 |
| Cellular responses to stress | 1.173 | -1.055 |
| C-type lectin receptors | 1.05 | -1.05 |
| Interleukin-1 family signaling | 1.127 | -1.013 |
| Immunoregulatory interactions between a Lymphoid and a non-Lymphoid cell | 0.953 | -0.939 |
| Fc epsilon receptor (FCERI) signaling | 0.976 | -0.928 |
| Metabolism | 0.933 | -0.913 |
| Extracellular matrix organization | 0.913 | -0.912 |
| Interleukin-17 signaling | 0.962 | -0.891 |
| Apoptosis | 0.772 | -0.649 |
| Signaling by VEGF | 0.892 | -0.5 |
| Complement cascade | 0.614 | -0.47 |
| Costimulation by the CD28 family | 0.541 | -0.419 |
| Regulation of TLR by endogenous ligand | 0.461 | 0.319 |
| Platelet homeostasis | 0.479 | 0.36 |
| Chemokine receptors bind chemokines | 1.171 | 0.52 |
| Interleukin-6 family signaling | 1.288 | 0.599 |
| Eicosanoid ligand-binding receptors | 0.731 | 0.684 |
| Interleukin-20 family signaling | 1.097 | 0.73 |
| Interleukin-2 family signaling | 1.057 | 0.767 |
| Interferon Signaling | 1.112 | 0.779 |
| Interleukin-12 family signaling | 1.475 | 0.923 |
| **Female PBS vs Female NP** | **Undirected global significance score** | **Directed global significance score** |
| Eicosanoid ligand-binding receptors | 1.787 | -1.535 |
| Metabolism | 1.458 | -1.005 |
| Platelet homeostasis | 1.71 | -0.917 |
| Regulation of TLR by endogenous ligand | 1.125 | -0.528 |
| Signaling by FGFR1 | 1.313 | -0.156 |
| Signaling by FGFR2 | 1.313 | -0.156 |
| Signaling by FGFR3 | 1.313 | -0.156 |
| Signaling by FGFR4 | 1.313 | -0.156 |
| C-type lectin receptors | 0.967 | 0.652 |
| Fc epsilon receptor (FCERI) signaling | 1.061 | 0.686 |
| Interleukin-17 signaling | 1.022 | 0.707 |
| Costimulation by the CD28 family | 0.776 | 0.72 |
| Interleukin-1 family signaling | 0.937 | 0.756 |
| Interleukin-20 family signaling | 0.871 | 0.859 |
| Complement cascade | 0.915 | 0.865 |
| Interleukin-6 family signaling | 0.867 | 0.867 |
| Interleukin-2 family signaling | 0.982 | 0.927 |
| Chemokine receptors bind chemokines | 0.962 | 0.962 |
| Interferon Signaling | 1.007 | 1.007 |
| Immunoregulatory interactions between a Lymphoid and a non-Lymphoid cell | 1.088 | 1.039 |
| Interleukin-12 family signaling | 1.059 | 1.059 |
| Cellular responses to stress | 1.341 | 1.063 |
| NF-κB pathway | 1.22 | 1.22 |
| Extracellular matrix organization | 1.776 | 1.697 |
| Apoptosis | 1.837 | 1.771 |
| Signaling by VEGF | 1.83 | 1.826 |
